# Mining the microbiota to identify gut commensals modulating neuroinflammation in a mouse model of multiple sclerosis

**DOI:** 10.1101/2021.11.10.468120

**Authors:** Paola Bianchimano, Graham J. Britton, David S. Wallach, Emma M. Smith, Laura M. Cox, Shirong Liu, Kacper Iwanowski, Howard L. Weiner, Jeremiah J. Faith, Jose C. Clemente, Stephanie K. Tankou

## Abstract

**Background:** The gut microbiome plays an important role in autoimmunity including multiple sclerosis and its mouse model called experimental autoimmune encephalomyelitis (EAE). Prior studies have demonstrated that the multiple sclerosis gut microbiota can contribute to disease hence making it a potential therapeutic target. In addition, antibiotic treatment has been shown to ameliorate disease in the EAE mouse model of multiple sclerosis. Yet, to this date, the mechanisms mediating these antibiotics effects are not understood. Furthermore, there is no consensus on the gut derived bacterial strains that drive neuroinflammation in multiple sclerosis.

**Results:** Here we characterized the gut microbiome of untreated and vancomycin treated EAE mice over time to identify bacteria with neuroimmunomodulatory potential. We observed alterations in the gut microbiota composition following EAE induction. We found that vancomycin treatment ameliorates EAE and that this protective effect is mediated via the microbiota. Notably, we observed increased abundance of bacteria known to be strong inducers of regulatory T cells, including members of Clostridium clusters XIVa and XVIII in vancomycin-treated mice during the presymptomatic phase of EAE, as well as at disease peak. We identified 50 bacterial taxa that correlate with EAE severity. Interestingly, several of these taxa exist in the human gut and some of them have been implicated in multiple sclerosis including *Anaerotruncus colihominis*, a butyrate producer, which had a positive correlation with disease severity. We found that *Anaerotruncus colihominis* ameliorates EAE and this is associated with induction of RORγt^+^ regulatory T cells in the mesenteric lymph nodes.

**Conclusions:** We identified vancomycin as a potent modulator of the gut-brain axis by promoting the proliferation of bacterial species that induce regulatory T cells. In addition, our findings reveal 50 gut commensals as regulator of the gut-brain axis that can be used to further characterize pathogenic and beneficial host-microbiota interactions in multiple sclerosis patients. Our findings suggest that elevated *Anaerotruncus colihominis* in multiple sclerosis patients may represent a protective mechanism associated with recovery from the disease.

## Background

Multiple sclerosis (MS) is a chronic immune mediated neurological disease characterized by infiltration of the central nervous system (CNS) with inflammatory leukocytes followed by demyelination and axonal loss[1–3]. An autoimmune response directed against components of myelin is the main pathogenic event during MS[4]. The exact autoimmune responses in MS are not fully known, although components of both the adaptive and innate immune systems are involved[5, 6]. Interferon-gamma (IFN-γ) producing Th1 and interleukin-17 (IL-17) secreting CD4^+^ T cells play a central role in the pathogenesis of MS[7, 8].

Genetic and environmental factors contribute to MS development[9]. Among the latter, several studies suggest that intestinal factors modulate MS disease severity[10]. In addition, alterations in the gut microbiota composition[11–16], gut derived products[17–19], intestinal permeability[20–22] and enteric nervous system functions[23] have been reported in MS patients. The human microbiome encompasses trillions of organisms that inhabit the gut and shape the gut-associated lymphoid tissue. Studies have shown that the gut microbiota shapes the development and function of both innate and adaptive immune cells[24–28]. Different commensals in the gut promote the differentiation of subsets of lymphocytes. Segmented filamentous bacteria (SFB) induces intestinal Th17[29], *Bacteroides fragilis* (*B. fragilis*) colonization of germ-free (GF) mice preferentially induces Th1 cells[30], and polysaccharide A of *B. fragilis* suppresses Th17 cells in conventional mice[31]. Clostridium clusters IV and XIVa promote T regulatory cells (Tregs) accumulation[32] and a subset of Clostridium clusters IV and XIVa which attach to the gut mucosa, can also promote Th17 cells[33].

The relationship between the host and its microbiota is generally mutually beneficial[34]. However, perturbations in the composition of the gut microbiota referred to as dysbiosis has been implicated with diseases of various etiologies, including autism, amyotrophic lateral sclerosis (ALS), Parkinson’s disease and several autoimmune disorders such as inflammatory bowel disease, diabetes, rheumatoid arthritis and MS[11, 12, 35–42]. Several studies have reported alterations in the gut microbiota composition of MS patients including increases in *Akkermansia muciniphila* and decreases in butyrate-producing bacteria[11, 14, 15, 40, 41, 43–46]. Whether these gut microbiota alterations in MS contribute to disease pathogenesis or are just a consequence remains unknown. We have previously reported that MS patients and mice at peak EAE have elevated intestinal miR-30d which increases the levels of *Akkermansia muciniphila* and ameliorates EAE[47]. Furthermore, we recently reported that MS-derived *A. muciniphila* attenuates EAE clinical scores[40]. These findings suggest that elevated *Akkermansia* in MS patients may be a consequence of the disease. The finding that transferring the gut microbiota from MS patients into mice exacerbates EAE, suggests that the MS gut microbiota can drive neuroinflammation[14, 15]. Consistent with these findings, others studies have reported that long-term antibiotic treatment in MS reduces relapse rates and gadolinium enhancing lesions as well as improves measures of disability[48, 49]. However, results across gut microbiota studies in MS lack consistency. Sample size, subject heterogeneity, study design, type of controls, geographical location, sequencing platforms and regions of 16S rRNA gene sequencing may all contribute to lack of reproducibility[50]. As such, the mechanisms through which altered gut-brain axis may contribute to CNS inflammation, demyelination and axonal loss remain poorly understood. This in turn makes it impossible to determine how the gut microbiome can be targeted for therapeutic purposes in MS.

Studies in animal models demonstrate that the gut microbiota can modulate neuroinflammation. Germ-free mice are protected from EAE and transferring specific pathogen free microbiota to these mice restore their susceptibility to EAE[51, 52]. Other studies have reported that oral treatment with broad-spectrun antibiotics significantly altered the gut microbiota and reduced EAE severity in a Treg-dependent manner[53]. Disease amelioration did not occur when antibiotics were given intraperitoneally, thereby bypassing the gut, suggesting that modulation of the gut microbiota produces protective effects. However these studies did not investigate the gut microbiome composition of the antibiotic treated mice and as such were not able to demonstrate that changes in the gut microbiota composition regulate CNS inflammation in EAE mice. Furthermore, SFB, a TH17 inducer, exacerbates EAE[52] whereas Tregs inducers such as polysaccharide A-positive *Bacteroides fragilis* ameliorates EAE[54]. Prior studies found decreased *Prevotella* in MS patients and mice fed human-derived *Prevotella histicola* are protected from EAE[11, 12, 15, 41, 55–57]. These findings suggest that bacteria depleted in MS can attenuate neuroinflammation. Furthermore, we reported that MS patients receiving a probiotic consisting of a mixture of *Lactobacillus-Bifidobacterium-Streptococcus* species displayed changes in their gut microbiota composition that were associated with anti-inflammatory immune markers in the periphery[58]. Another study reported that supplementation of MS patients with a mixture of three *Lactobacillus* species and one *Bifidobacterium* species was associated with improved EDSS score as well as decreased depression and stress[59]. Taken together, these studies demonstrate that modulating the microbiota has therapeutic potential and the EAE mouse model is a useful tool to identify human gut derived bacteria with neuroimmunomodulatory potential.

In this report, we investigated the gut microbiota of untreated and vancomycin-treated EAE mice at multiple time points spanning pre- and post-immunization state with myelin oligodendrocyte glycoprotein (MOG) to identify bacterial taxa that correlate with EAE severity and as such have the potential to modulate CNS inflammation. We found that vancomycin treatment ameliorates EAE and that this protective effect is mediated via the gut microbiota. Notably, we observed increased abundance of Tregs inducing bacteria including members of Clostridium clusters XIVa and XVIII in the gut of vancomycin treated mice compare to untreated mice. Importantly, many of the bacteria that correlated with EAE severity exist in the human gut microbiome. In addition, several of the bacteria that have been implicated in MS such *A. muciniphila* and *Anaerotruncus colihominis* correlated with EAE severity. We found that human derived *A. colihominis* ameliorates EAE and this is associated with the induction of RORγt^+^ regulatory T cells in the mesenteric lymph nodes. Our work led to the identification of 50 bacterial species with neuroimmunomodulatory potential. This list of bacteria can serve as a starting point to define pathogenic and beneficial host-microbiota interactions that modulate neuroinflammation in MS. This information will in turn facilitate the development of innovative microbiota-based approaches to target neuroinflammation in MS.

## Materials and methods

### Mice

C57BL/6J female mice were obtained from The Jackson Laboratory and kept in a specific pathogen-free facility at the Harvard Institute of Medicine or Icahn School of Medicine at Mount Sinai on a 12-hour light/dark cycle. C57BL/6J germ-free female mice were obtained from the Massachussetts Host-Microbiome Center at the Brigham & Women’s Hospital and kept in a specific pathogen-free facility at the Harvard Microbiome Facility or the Massachussetts Host-Microbiome Center at the Brigham & Women’s Hospital. Mice were all 8-10 weeks of age and cohoused, four mice from same experimental condition per cage. Mice were assigned randomly to the experimental groups. Mice were fed an ad-libitum diet of Picolab Rodent Diet 5053 and distilled water without added preservatives (provided by animal facility). Animals were housed in a biosafety level 2 facility using autoclaved cages and aseptic handling procedures and kept under a 12-hour light/dark cycle. All animal experiments described in this paper were approved by the institutional Animal Care and Use Committee (IACUC) at Harvard Medical School or Icahn School of Medicine at Mount Sinai and carried out in accordance with those approved animal experiment guidelines.

### EAE Induction

EAE was induced by injecting 8 to 10-week-old female C57BL/6J mice with 150 mg MOG_35-55_ peptide (Genemed Synthesis) emulsified in complete Freund’s adjuvant (CFA) (BD Difco) per mouse subcutaneously in the flanks, followed by intraperitoneal administration of 150 ng pertussis toxin (List biological laboratories, Inc.) per mouse on days 0 and 2 as previously described[60]. Clinical signs of EAE were assessed according to the following score: 0, no signs of disease; 1, loss of tone in the tail; 2, hind limb paresis; 3, hind limb paralysis; 4, tetraplegia; 5, moribund. Differences between the groups were determined by Friedman test and Dunn correction for multiple comparisons.

### Isolation and Identification of *Enterococcus faecalis*

Adult C57BL6/J female mice received vancomycin (0.5g/L) in drinking water for 2 weeks. Next, mice were euthanized, cecum was rapidly collected and resuspended in prereduced anaerobically sterilized saline and 100uL of 10^−4^ through 10^−7^ dilutions was plated on brucella blood agar (Anaerobe Systems) and incubated in an aerobic incubator. Eight to ten colonies were isolated in pure culture after 48 hrs incubation. Pure *Enterococccus faecalis* cultures were identify by biotyping using the Bruker Biotyper LT MALDI-TOF mass spectrometer. The *E. faecalis* ID was confirmed by amplifying the nearly full-length 16S rRNA gene using the 8F and 1510R primers according to previous methods[61]. After polymerase chain reaction, PCR products were sequenced by Sanger Sequencing at Genewiz. Identification and percent identity were then performed using batch BLAST, National Centers for Biotechnology Information.

### Bacteria strains, Growth and Administration

*Anaerotruncus colihominis* (DSMZ, DSM#:17241) and *Enterococcus faecalis* were grown anaerobically at 37°C in Brain Heart Infusion (BHI) medium (Cat#: DF0037178; Fisher Scientific). Each bacterial culture suspension was subsequently transferred to 3 Brucella Blood agar plates and colonies from all plates were resuspended in 2.5mL anaerobic sterile PBS yielding a suspension of live bacteria at a density of OD600= 1.3. 8-week-old female C57BL/6J mice were colonized with 200uL of bacteria suspension via oral gavage 3 days per week beginning 3 weeks prior to disease induction and bacteria treatment was received for the entire duration of the experiment. Control mice were orally gavaged with anaerobic PBS (vehicle).

### Histopathology

Mice were euthanized at the termination of experiments and were intracardially perfused with PBS. Lumbosacral spinal cords were fixed with Formalin. Tissue was processed and stained as previously described[60]. Paraffin embedded serial sections were stained with Luxol Fast Blue for myelin, Bielschowsky silver for axons. The demyelinated area and axonal/neuronal loss were determined using ImageJ software (National Institutes of Health, USA) and the percentages of demyelinated and axonal/neuronal lost area out of total area were calculated. To detect immune infiltrate, spinal cord sections were stained using Rat anti mouse CD4 antibody (1:100; eBioscience, Cat# 14-0042-82) with secondary biotinylated antibodies. Avidin-peroxidase and 3,4-Diaminobenzidine was used as the color substrate. CD4+ cells were counted manually using multi point function on ImageJ, and cell density calculated as number of cells per total white matter area.

### Antibiotic Treatment

In order to investigate the effect of vancomycin and neomycin on EAE development, mice were given vancomycin 0.5mg/mL or neomycin 1mg/mL (Fisher Scientific) in drinking water for 2 weeks (Figure 2A). For the cohousing experiment, mice received vancomycin 3mg in 200uL nuclease free water via oral gavage (Figure 2E). For the fecal transfer experiment using wild type C57BL/6J mice, to deplete bacteria, mice were given a mixture of antibiotics (ampicillin 1 mg/mL, vancomycin 0.5 mg/mL, neomycin 1 mg/mL, metronidazole 1 mg/mL; Fisher Scientific) in drinking water for 3 consecutive days (Figure 2G).

### Fecal Microbiota Transplantation

For mice cecal transfer experiment, 1 mouse cecum was homogenized in 8mL of sterile anaerobic PBS and 200uL of the cecal slurry was administered to recipient adult conventionally raised female C57BL/6J mice or 4-week-old female C57BL/6J germ free mice by oral gavage at the indicated times in the figures.

### 16S rRNA gene sequencing and analysis

Feces were collected at 11 time points spanning pre- and post-immunization states: before vancomycin treatment (day -22), on vancomycin (days -21, -20, -18, -14), at vancomycin discontinuation (day -6), post vancomycin discontinuation/pre-EAE induction (day 0), post EAE induction-latent period (days 3, 8), post EAE induction-peak disease (day 15), post EAE induction-recovery phase (day 29). Fecal samples were collected immediately upon defecation, snap frozen and kept at -80C until processed. Bacterial DNA was isolated from feces using MoBio PowerLyzer PowerSoil Kit (Qiagen). Amplicons spanning variable region 4 (V4) of the bacterial 16S rRNA gene were generated with primers containing barcodes (515F, 806R) from the Earth Microbiome project[62] using HotMaster Taq and HotMaster Mix (QuantaBio) and paired-end sequenced on an Illumina MiSeq platform at the Harvard Medical School Biopolymer Facility. Paired-end 16S rRNA gene reads were trimmed for quality (target error rate < 0.5%) and length (minimum 125bp) using Trimmomatic, merged using FLASH (Fast Length Adjustment of Short reads), and quality screened using QIIME 1 (Quantitative Insights Into Microbial Ecology). Spurious hits to the PhiX control genome were identified using BLASTN (Basic Local Alignment Search Tool) and removed. Passing sequences were trimmed of primers, evaluated for chimeras and screened for mouse-associated contaminant using Bowtie2. True 16S rRNA sequence fragments were confirmed as such by requiring at least one BLAST match against the GreenGenes database (default parameters). Greengenes was only used for this QC step, and not for taxonomic assignment. Chloroplast and mitochondrial contaminants were detected and filtered using the RDP (Ribosomal Database Project) classifier with a confidence threshold of 80%. High-quality 16S rRNA amplicon sequences were assigned to a high-resolution taxonomic lineage using Resphera Insight (v.2.2), for which the performance has been previously benchmarked using high-quality draft genome assemblies of well-defined species from the Human Microbiome Reference Genome Database (http://hmpdacc.org/HMRGD/; ref. 13). This method utilizes a manually curated 16S rRNA database of 11,000 unique species, and a hybrid global-local alignment procedure to assign short next-generation sequencing sequences from any region of the 16S rRNA gene to a high-resolution taxonomic lineage[63]. Operational taxonomic units (OTUs) assignment was performed as follows: sequences that could not be confidently assigned to any known species were clustered de novo and clusters were given an OTU number identifier and annotated with a closest relative or set of relatives as previously described[64]. Taxonomy summary plots were generated based on relative abundances as implemented in QIIME. Testing for significant differences in alpha diversity was performed by ONE WAY ANOVA followed by Tukey’s post-hoc test or a Mixed-Effects Model when comparing matched data with a missing value followed by Dunnett’s corrections (as indicated on the figure legends). Beta diversity was estimated using Bray-Curtis, and distances then used to perform Principal Coordinate Analysis as implemented in QIIME. Differences in beta diversity were tested using PERMANOVA with FDR correction for multiple comparison testing. Compositional differences were determined using linear discriminant analysis effect size (LEfSe) with alpha set at 0.05 and the effect size set at greater than 2. To identify bacteria linked with EAE severity, Spearman correlations were performed in R using the function Cor.test and FDR adjustment was performed in R using the function p.adjust.

### Fecal Microbe Quantification by qPCR

DNA extracted from feces as described above was used for specific bacteria abundance. Quantitative PCR (qPCR) analysis was conducted using a QuantStudio 7 Flex Real-Time PCR System (Applied Biosystems). *A. colihominis, E. faecalis and SFB* were quantified by SYBR Green (Applied Biosystems), and primer pairs as follows: Allbacteria (universal 16S rRNA gene, reference): Forward: ACTCCTACGGGAGGCAGCAGT, Reverse: ATTACCGCGGCTGCTGGC[65]; *A. colihominis* 16S rRNA gene: Forward: GGAGCTTACGTTTTGAAGTTTTC, Reverse: CTGCTGCCTCCCGTA [66]; *E. faecalis* 16S rRNA gene: Forward: TACTGACAAACCATTCATGATG, Reverse: AACTTCGTCACCAACGCGAAC[67]; *SFB* 16S rRNA gene: Forward: GACGCTGAGGCATGAGAGCAT, Reverse: GACGGCACGGATTGTTATTCA[65]. The relative quantity was calculated using the comparative CT method normalizing to the amount of all bacteria in the sample[68].

### Immune cell isolation

For immune profiling, naïve 8-week-old female C57BL/6J mice were orally gavaged with *A. colihominis* or *E. faecalis* or PBS 3 days per week for 3 weeks as described above after which mice were euthanized using carbon dioxide and tissues rapidly dissected. Single cell suspensions were obtained as previously described[69]. Mesenteric lymph nodes and spleens were dissociated by pressing through 70 um mesh and red blood cells removed from splenocytes using ALK lysis buffer. Intestinal tissues were cleaned of feces, Peyer’s patches removed and deepithelialized in 5mM buffered EDTA before digestion with 0.5mg/ml Collagenase Type IV (Sigma C5138) and 0.5 mg/ml DNase1 (Sigma DN25). Cell suspensions were filtered through 40 um strainers and washed before use. No further enrichment of lymphocytes was performed.

### Flow cytometry

For analysis of cytokine production, mononuclear cell suspensions were restimulated with 5 ng/mL phorbal 12-myristate 13-acetate (PMA), 500 ng/mL ionomycin with monensin (Biolegend) for 3.5 hours at 37°C. All other analysis was performed on unstimulated cells. Intracellular cytokine staining was performed following surface staining and fixation with IC Fixation Buffer (ThermoFisher/eBioscience). Transcription factor staining was performed using FoxP3 Fixation/Permeabilization buffers (ThermoFisher/eBioscience). Super Bright Complete staining buffer (ThermoFisher) was included when multiple Brilliant Violet-conjugated antibodies were used together. The following anti-mouse antigen fluorochrome-conjugated antibodies were used, obtained from BioLegend unless otherwise stated: CD45 Brilliant Violet 750 (30-F11), CD45 APC-Cy7 (30-F11), CD4 FITC (RM4-5), CD4 PerCp-Cy5.5 (RM4-5), CD3 AlexaFluor 700 (17A2), CD3 PerCP-eFluor710 (17A2, ThermoFisher/eBioscience), CD8-alpha Pacific Orange (5H10, ThermoFisher), CD11b PerCP-Cy5.5 (M1/70), CD11c PE-Cy7 (N418), CD103 Brilliant Violet 510 (2E7), CD64 Brilliant Violet 786 (X54-5/7.1, BD Bioscience), MHC-II I-A/I-E Pacific Blue (M5/114.15.2), CD80 Brilliant Violet 421 (16-10A1), CD86 Brilliant Violet 605 (GL-1), Ly-6G Brilliant Violet 570 (1A8), FoxP3 PE (FJK-16s, ThermoFisher/eBioscience), RORγt APC (B2D, BD BioScience), GATA3 Brilliant Violet 421 (16E10A23), IL-17A PE (TC11-18H10.1), IL-10 APC (JES5-16E3), IFNγ PE-Cy7 (XMG1.2), GM-CSF FITC (MP1-22E9). Dead cells were excluded using Zombie Aqua (BioLegend) or eFluor780 Fixable Viability Dye (ThermoFisher/eBioscience). Data was acquired on a five laser Aurora Cytometer (Cytek Biosciences) and raw data was spectrally unmixed using SpectroFlo software (Cytek Biosciences). Unmixed data files were analyzed using FlowJo 10 (BD Biosciences) and statistical analyses performed using R Studio 1.1.463 and Prism 6 (Graphpad).

### Quantification and Statistical Analysis

All graphs, calculations, and statistical analyses were performed using the GraphPad Prism software for Mac (GraphPad Software, San Diego, CA, USA). Comparisons of three or more groups following a normal distribution were performed using one-way ANOVA followed by Dunnett’s test or Tukey’s test as specified in figures. For datasets of non-normal distributions, the Mann Whitney test was used for comparisons between two groups. EAE clinical scores were analyzed over time with the non-parametric Friedman test for repeated measurements as specified in figures[70, 71]. Exact statistical instruments, sample sizes, and P values are indicated in each figure. We did not use statistical methods to determine sample size; we used sample sizes that were similar to those in our previous publications andthose of others[60, 72–75].

## Results

### Characterization of the gut microbiome during EAE

To investigate changes in the gut microbiota composition during EAE, we collected fecal pellets at 11 time points spanning pre- and post-MOG immunization state from 8-week-old C57BL/6J (B6) mice (**Fig. 1A, 1B**). Next, we analyzed the microbiota from these mice by sequencing the V4 region of the microbial 16S rRNA gene. We first assessed alpha diversity and found no significant changes at any time point (**Fig 1C, S1A**). We then examined beta diversity to assess whether the composition of the microbiota changes during EAE and found no significant differences in community structure between pre- and post-immunization states (**Fig. 1D**). We next investigated whether the relative abundances of bacteria differed between pre- and post-immunization states at the species level. We observed a change in the abundance of species belonging to several genera including *Akkermansia, Turicibacter, Lactobacilli* and *Clostridium* (**Fig. 1E, S1B**). We observed that most changes in taxa abundance occurred within the first 2 weeks post EAE induction (**Fig. 1F, S2**). In particular, we found increased levels of *Clostridium cocleatum, Ruminococccus flavefaciens, Clostridium ruminantium* and *Clostridium chauvoei* at 3 days post immunization (DPI) compared to 0 DPI (**Fig. 1F, S2**). We observed increased abundance of 8 taxa including *Clostridium scindens*, a member of Clostridium cluster XIVa as well as three bacteria belonging to Clostridium sensu stricto: *C. chauvoei/quinii* operational taxonomic unit (OTU) 8778, *C. chauvoei* and *C. celatum* at 8 DPI compared to 0 DPI (**Fig. 1F-G, S2**). Family *Verrucomicrobiaceae* which contains species *Akkermansia muciniphila* were enriched in the gut of mice at 15 DPI compared to 0 DPI. These findings are consistent with prior studies reporting increased *A. muciniphila* in EAE mice [47, 76, 77]. In addition, several studies have reported increased level of *A. muciniphila* in MS compare to healthy control[11, 14, 15, 40, 46]. *C. scindens* was also increased at 15 DPI compared to 0 DPI (**Fig. 1F-G, S2**). Interestingly, we have recently reported that *C. scindens* is increased in MS patients compare to healthy control[40]. Two *Clostridia* species were decreased at 3 DPI compared to 0 DPI: *Clostridium viride* (OTU 5938) a member of Clostridium cluster IV, and *Clostridium indolis* (OTU 1977), a member of Clostridium cluster XIVa (**Fig. 1F-G, S2**). Furthermore, several species belonging to Clostridium cluster XIVa were decreased at 15 DPI compared to 0 DPI: *C. boltea/clostridiforme/oroticum* (OTU 8033), *C. polysaccharolyticum* (OTU 7948), *C. indolis* (OTU 1977), *C. hylemonae* and *C. aldenense/indolis* (**Fig. 1F-G, S2**). These results are consistent with findings from one study reporting decreased abundance of species belonging to Clostridium cluster IV and XIVa in MS patients[41]. We also found decreased *Dorea formicigenerans* (OTU 2094) and *Lactobacillus gasseri/hominis/johnsonii/taiwanensis* at 3 DPI compared to 0 DPI (**Fig. 1F-G, S2**). These findings are consistent with results from prior studies reporting decreased genera *Dorea* and *Lactobacilli* as well as decreased *L. johnsonii/taiwanensis/gasseri* in EAE mice[76, 78]. *Turicibacter sanguinis, L. gasseri/hominis/johnsonii/taiwanensis* and *Olsenella profusa* (OTU 3942) were also decreased at 15 DPI compared to 0 DPI (**Fig. 1F-G, S2**).

**Figure 1.**
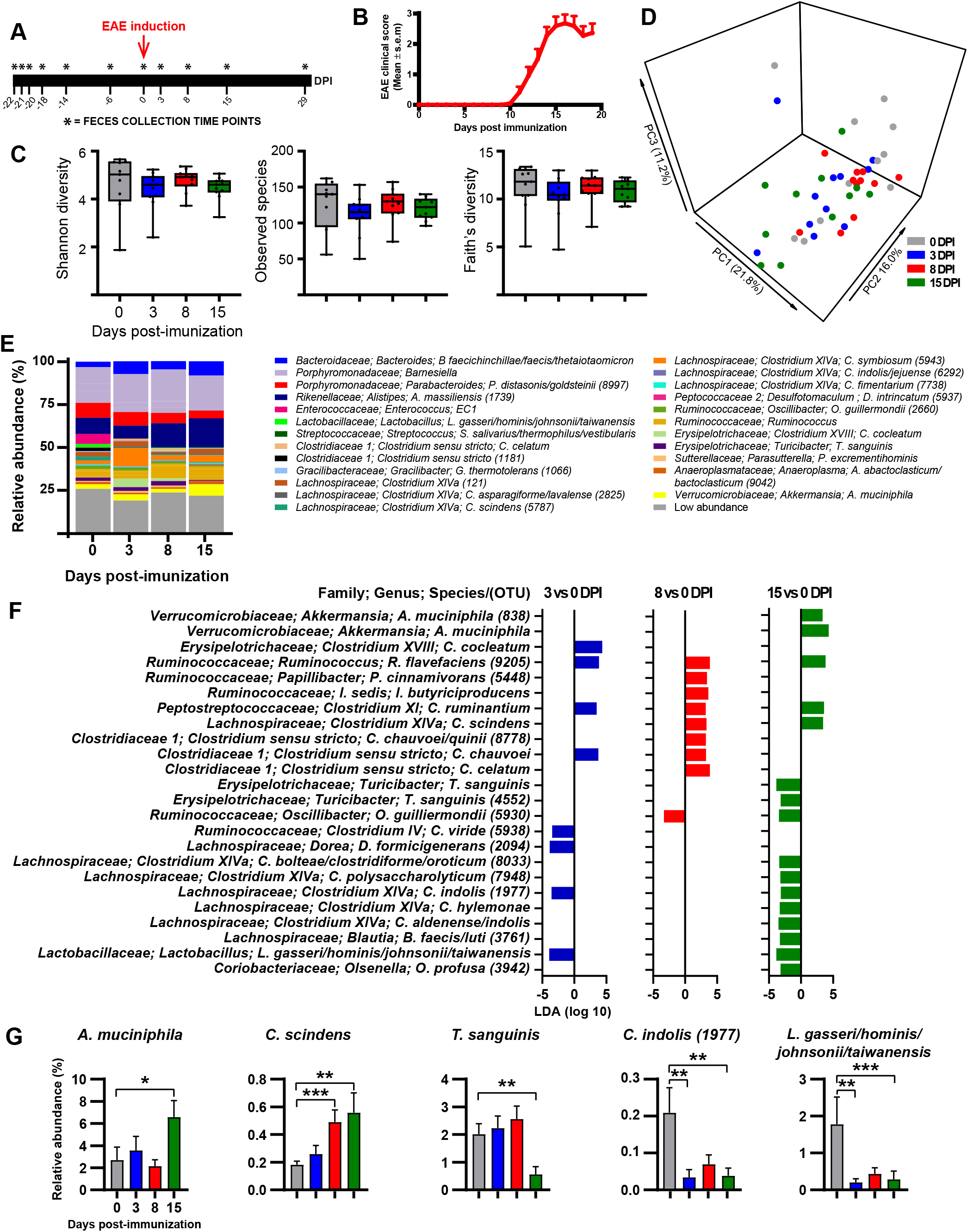
Changes in microbiota composition during EAE. Mice were immunized with MOG and feces were collected prior to EAE induction at -22, -21, -20, -18, -14 and -6 days post immunization (DPI), on the day of immunization prior to EAE induction at 0 DPI and post EAE induction at 3, 8, 15 and 29 DPI. **A**. Schematic design showing the 11 time points for fecal samples collection. **B**. Mean EAE clinical scores overtime. Results are presented as Mean ± SEM (n= 11 mice). **C**. α-Diversity metrics for Shannon diversity, Observed species and Faith’s Diversity were calculated at an average sampling depth of 1,500 reads per sample. No significant differences were observed for any of the diversity estimators analyzed (Mixed-effect model followed by Dunnett’s test) **D**. Principal coordinate analysis of intestinal microbiota samples based on Bray Curtis. Each dot represents the microbiota from one mouse. **E**. Taxa plots showing compositional differences in fecal microbiota in mice at the indicated time points. **F**. Linear discriminant analysis (LDA) effect size of significantly altered bacteria at the lowest classifiable levels at the indicated time points. **G**. Relative abundance of selected species altered in EAE mice at the indicated time points. Results are presented as mean ± SEM (n = 11 mice), *p<0.05; **p<0.01; ***p<0.001. OTU = Operational taxonomic unit; Numbers in parenthesis represent OTU. EC1: *Enterococcus canintestini/canis/dispar/ durans/faecalis/faecium/hirae/ lactis/mundtii/ratti/rivorum/villorum*.

### Amelioration of EAE after vancomycin is mediated via the microbiota

To this date, it remains unclear if the neuroprotective effects of antibiotics in EAE mice are driven by changes in the gut microbiota composition. To investigate this, 8-week-old B6 mice were treated with vancomycin or neomycin, two poorly absorbable antibiotics, for two weeks, after which antibiotics were discontinued (**Fig. 2A**). Next, mice were immunized with MOG for EAE induction. We found that, compared to control mice, vancomycin-treated mice had significantly less severe disease, whereas neomycin-treated mice behaved like the control group (**Fig. 2B-D**). Hence, these results are consistent with our prior report that vancomycin treatment ameliorates EAE[78]. To determine if the protective effect of vancomycin is mediated via the microbiota, we conducted three follow-up experiments. First, we did a cohousing experiment in which 8-week-old B6 mice receiving normal drinking water or vancomycin once a day via oral gavage were either housed with the same treatment group (single treatment) or co-housed between treatment groups. Two weeks later, vancomycin was discontinued and mice were immunized with MOG to induce EAE (**Fig. 2E**). As expected, single-treatment housed control mice developed more severe disease than the single-treatment housed vancomycin mice (**Fig. 2F**). We observed that vancomycin treated mice that were co-housed with control mice developed more severe disease than single-treatment housed vancomycin mice (**Fig. 2F**). We also found that control mice that were co-housed with vancomycin treated mice had delayed disease onset compared to single-treatment housed control mice (**Fig. 2F**). Second, we performed a cecal transfer experiment where 8-week-old conventionally raised B6 mice were treated with an antibiotic mixture containing neomycin, vancomycin, ampicillin and metronidazole for three days. Twenty-four hours after antibiotics discontinuation, half of these mice were fed feces from control mice and the remaining half received feces from vancomycin treated mice. Ten days post gavage, EAE was induced and EAE scores were monitored overtime (**Fig. 2G**). We found that mice that received feces from vancomycin treated mice had less severe disease than mice who were fed feces from control mice (**Fig. 2H**). Third, we conducted a second cecal transfer experiment, this time using 4-week-old B6 germ-free (GF) mice that were either gavaged with feces from control mice or with feces from vancomycin treated mice. 4-week post gavage, EAE was induced and EAE scores were monitored overtime (**Fig. 2I**). We observed that mice that were fed feces from vancomycin treated mice had less severe EAE compared to those that received feces from control mice (**Fig. 2J-L**). Taken together, these results suggest that the vancomycin protective effect is mediated via the microbiota. Segmented filamentous bacteria (SFB) exacerbates EAE via induction of Th17 cells and given that SFB is sensitive to vancomycin, we repeated the fecal transfer experiment in GF mice using wild type B6 mice microbiota from the Jackson laboratory which has been previously shown to be free of SFB[29]. We confirmed that the microbiota from our B6 mice lacks SFB by quantitative real time PCR using SFB specific primers (**Fig. S3A-B**). Next, 4-week-old GF mice were gavaged with SFB free microbiota and 3 weeks later half of the mice were treated with vancomycin via oral gavage once daily for 2 weeks (**Fig. S3C**). We found that vancomycin treatment protects mice from EAE even in the absence of SFB (**Fig. S3D-F**). Hence, the protective effect of vancomycin is not modulated by SFB.

**Figure 2.**
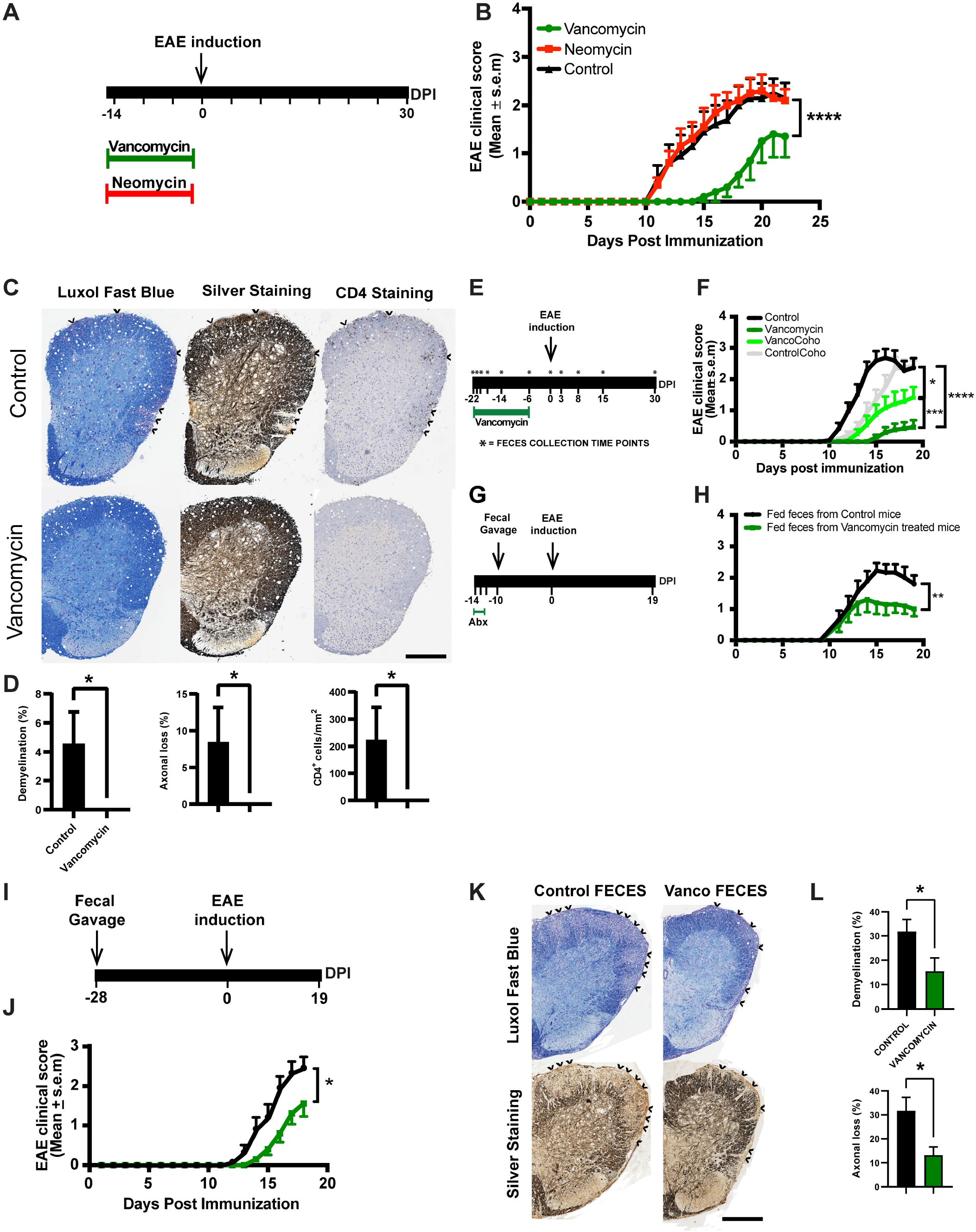
Effect of vancomycin on EAE development. **A**. Schematic representation showing the time course for Vancomycin//Neomycin treatments. **B**. Mean EAE clinical scores overtime. Results are presented as mean ± SEM (n= 10 mice/group). ****p<0.0001, Freidman test with Dunn correction for multiple comparisons. **C**. Histopathological evaluation of demyelination with Luxol Fast Blue (LFB), axonal loss with Bielschowsky’s silver (silver) staining and CD4^+^ cell infiltrates at 15 DPI. Arrows denote demyelination (LFB), axonal loss (silver staining) and CD4^+^ cell infiltrates of representative spinal cord sections from control and vancomycin treated EAE mice. Scale bars, 500µm. **D**. Quantification of demyelination, axonal loss and CD4^+^ cell infiltrates of individual mouse. Representative data of three independent experiments with n=5 mice per group are shown. Error bars denote mean ± SEM; Mann Whitney test was performed. *p<0.05. **E**. Experimental scheme of cohousing experiment. Conventionally raised mice were either untreated or treated with vancomycin daily via oral gavage for 2 weeks. Mice were either housed with the same treatment group (single treatment) or cohoused between treatment groups. Mice were immunized with MOG 1 week post discontinuation of vancomycin for EAE induction. **F**. Mean EAE clinical scores overtime in single-treatment housed untreated mice (Control), single-treatment housed vancomycin mice (Vancomycin), untreated mice cohoused with vancomycin treated mice (ControlCoho) and vancomycin treated mice cohoused with untreated mice (VancoCoho). Error bars denote Mean ± SEM (n= 11 mice/group); the Friedman test based on scores from 0 DPI until the end of the experiment and Dunn’s multiple-comparison test were performed. *p<0.05; ***p<0.001; ****p<0.0001. **G**. Experimental scheme of fecal transfer experiment in conventionally raised mice. Conventionally raised mice were on drinking water supplemented with ampicillin (1g/L), neomycin sulfate (1g/L), metronidazole (1g/L) and vancomycin (0.5g/L) for 3 days. 48 hrs post antibiotics discontinuation, half of the mice were orally gavage with feces from wild type B6 mice and the remaining half was fed feces from vancomycin-treated mice 10 days prior to EAE induction. Mice were immunized with MOG for EAE induction. **H**. Mean EAE clinical scores overtime. Error bars denote Mean ± SEM (n= 10 mice/group); the Friedman test based on scores from 0 DPI until the end of the experiment and Dunn’s multiple-comparison test were performed. **p<0.01 **I**. Experimental scheme of fecal transfer experiment in germ-free mice. Germ-free mice were either fed feces from untreated or vancomycin-treated coventionally-raised mice. 4 weeks post oral gavage, colonized germ-free mice were immunized with MOG for EAE induction. **J**. Mean EAE clinical scores overtime. Error bars denote Mean ± SEM (n=10-12 mice/group); the Friedman test based on scores from 0 DPI until the end of the experiment and Dunn’s multiple-comparison test were performed. *p<0.05. **K**. Histopathological evaluation of demyelination with Luxol Fast Blue (LFB) and axonal loss with Bielschowsky’s silver (silver) staining. Arrows denote demyelination (LFB) and axonal loss (silver) staining of representative spinal cord sections. Scale bars, 500µm. **L**. Quantification of demyelination and axonal loss of individual mouse. Representative data of three independent experiments with n=5 mice per group are shown. Error bars denote mean ± SEM; Mann Whitney test was performed. *p<0.05.

### Vancomycin effect on the gut microbiota during EAE

Given that vancomycin treated mice had less severe EAE and we found that this protective effect is mediated via the microbiota, we next profiled the 16S rRNA gene of these mice. Examining alpha diversity, as expected, we found that vancomycin treated mice had decreased Shannon diversity, Observed species and Faith’s diversity compared to control mice (**Fig. 3A**). However, starting at 3 DPI, we found no change in alpha diversity between vancomycin treated mice that were co-housed with control mice and single-treatment housed control mice (**Fig. 3A**). In addition, starting at 0 DPI, co-housed vancomycin treated mice had increased Shannon diversity compared to single-treatment housed vancomycin mice (**Fig. 3A**). We also observed increased Observed species and Faith’s diversity in co-housed vancomycin treated mice compare to single-treatment housed vancomycin mice starting at 0 DPI (**Fig. 3A**). Examining beta diversity, the gut microbiota composition was not significantly different among the 4 mice groups at -22 DPI (baseline) prior to vancomycin initiation (q>0.29). We observed that overall microbiota community structure of single-treatment housed vancomycin mice differed from that of single-treatment housed control mice (q<0.003**; Fig. 3B**) as well as that of cohoused control mice (q<0.002**; Fig. 3B**) starting at -21 DPI (one day post vancomycin initiation). At -21 DPI, singly housed vancomycin treated mice clustered with co-housed vancomycin mice (q=0.192**; Fig. 3B**). However, starting at -6DPI, single-treatment housed vancomycin mice clustered differently from cohoused vancomycin mice (q<0.002**; Fig. 3B**). Hence, despite receiving the same vancomycin treatment, single-treatment housed vancomycin mice and cohoused vancomycin treated mice have different alpha diversity and microbial composition. In addition, the overall microbiota community structure differed between cohoused vancomycin treated mice and single-treatment housed control mice at all time points (q<0.002**; Fig. 3B**) and between cohoused vancomycin treated mice and cohoused control mice at -21, -6 and 0 DPI (q<0.005; **Fig. 3B**). However, starting at 3 DPI, cohoused vancomycin treated mice clustered with cohoused control mice (q>0.05**; Fig. 3B**).

**Figure 3.**
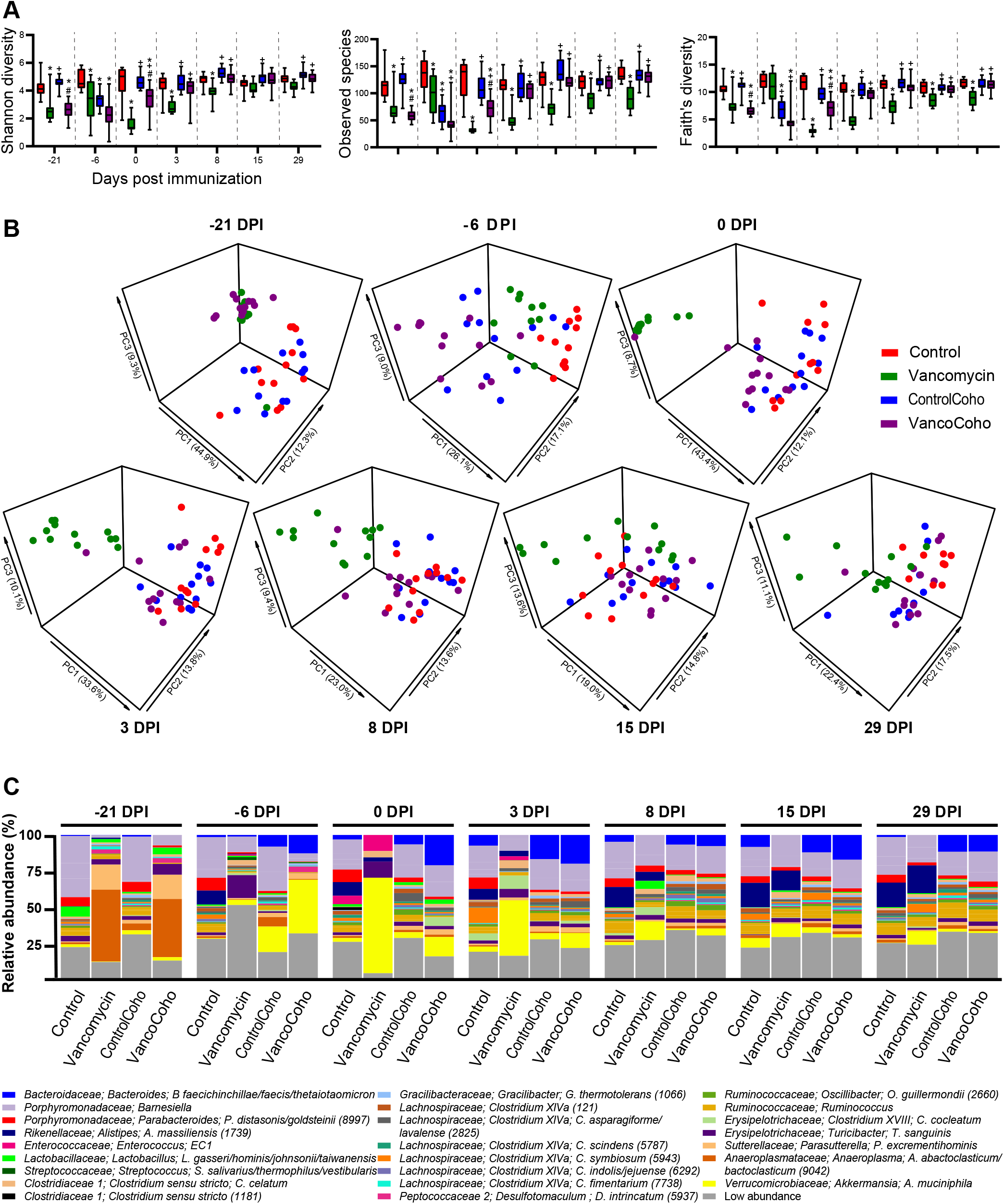
Effect of vancomycin on Alpha and Beta-diversity during EAE. **A**. Alpha diversity metrics for Shannon diversity, Observed species and Faith’s diversity were calculated at an average sampling depth of 1,500 reads per sample in single-treatment housed untreated mice (Control), single-treatment housed vancomycin mice (Vancomycin), untreated mice cohoused with vancomycin treated mice (ControlCoho) and vancomycin treated mice cohoused with untreated mice (VancoCoho). Results are expressed as mean ± SEM (n = 11-12 mice/group). Statistical comparisons are made using one-way ANOVA followed by Tukey test. * = significant difference with control group (p<0.05); + = significant difference with vancomycin group (p<0.05); # = significant difference with control-cohoused group (p<0.05). **B**. Principal coordinate analysis of intestinal microbiota samples based on Bray Curtis show significantly different clustering between control and vancomycin (q<0.003), Controlcoho and Vancomycin (q<0.002), Vancocoho and Control (q<0.002) at all time points, Vancomycin and Vancocoho (q<0.002), Control and Controlcoho (q<0.002) starting at -6 DPI, Vancocoho and Controlcoho (q<0.005) at -21, -6 and 0 DPI but not at 3,8,15 and 29 DPI. q= PERMANOVA p values adjusted for false discovery rate. Each dot represents the microbiota from one mouse. **C**. Taxa plots showing compositional differences in fecal microbiota at the indicated time points in Control, Vancomycin, ControlCoho and VancoCoho mice. EC1: *Enterococcus canintestini/canis/dispar/ durans/faecalis/faecium/hirae/ lactis/mundtii/ratti/rivorum/villorum*.

To identify taxa that account for the difference in beta diversity in these mice, we generated a taxa plot. As expected, we observed a change in the abundance of several bacteria between vancomycin treated and untreated mice including *A. muciniphila, T. sanguinis, Lactobacilli, Enterococcus* and *Clostridium* species (**Fig. 3C**). We found several taxa that are increased in single-treatment housed vancomycin mice compared to single-treatment housed control mice or cohoused vancomycin treated mice including: *A. muciniphila, T. sanguinis, Lactobacillus gasseri/hominis/johnsonii/taiwanensis* as well as several *Clostridium* species belonging to Clostridium cluster XIVa and Clostridium sensu stricto (**Fig. 4A-B, S4A-B**). These findings are consistent with our previous report that *A. muciniphila* ameliorates EAE by inducing Tregs[47] as well as several studies observing that administering various *Lactobacilli* strains to mice leads to an attenuation of EAE clinical course by either inducing Tregs or suppressing Th17 cells induction[79–81]. We also found that single-treatment housed vancomycin mice had decreased level of most species belonging to Clostridium cluster XIVa that we were able to detect during the pre-symptomatic stages of EAE (0, 3 and 8 DPI) (**Fig. S4A**). However, at peak disease (15 DPI), we observed an increase of four species belonging to Clostridium cluster XIVa in single-treatment housed vancomycin mice compare to single-treatment housed control mice including *Clostridium scindens* (OTU 2213), *Clostridium asparagiforme/lavalense* (OTUs 1808 and 2825) and *Clostridium indolis/jejuense* (OTU 6292) (**Fig. 4A, S4A**). In addition, *Clostridium cocleatum*, a member of Clostridium cluster XVIII, was increased in single-treatment housed vancomycin mice compare to single-treatment housed control mice or cohoused vancomycin mice (**Fig. 4A-B, S4A-B**). Interestingly, species that fall within clusters XIVa and XVIII of *Clostridia* are strong Tregs inducers[32, 82, 83]. When comparing the microbiota composition of vancomycin treated mice and control mice that were cohoused, *A. muciniphila* was increased in cohoused vancomycin treated mice compare to cohoused control mice (**Fig. 4C, S4C**). We also observed that *C. indolis/jejuense* (OTU 6292) and *C. oroticum* (OTU 2322), members of Clostridium cluster XIVa, as well as *C. cocleatum* were increased in cohoused vancomycin treated mice compare to cohoused control mice (**Fig. 4C, S4C**). Furthermore, *Coprobacillus cateniformis* (OTU 4628), a butyrate producer[84], was increased in cohoused vancomycin treated mice compare to cohoused control mice (**Fig. 4C, S4C**). Interestingly, butyrate is a Tregs inducer that has been shown to ameliorate EAE[85, 86]. Taken together, these findings suggest that a potential mechanism by which vancomycin treatment attenuates EAE severity is via promoting the proliferation of bacterial species that are Tregs inducers. These results also suggest that single-treatment housed vancomycin mice had less severe disease than cohoused vancomycin treated mice because they had more Tregs inducing bacterial species. Several bacteria were decreased in single-treatment housed vancomycin mice compared to single-treatment housed control mice or cohoused vancomycin treated mice. Notably, *Anaerotruncus colihominis* (OTU 2694) was depleted in single-treatment housed vancomycin mice (**Fig. 4A-B, S4A-B**). Interestingly, we and others have previously reported increased *A. colihominis* in MS patients compare to healthy subjects[19, 40].

**Figure 4.**
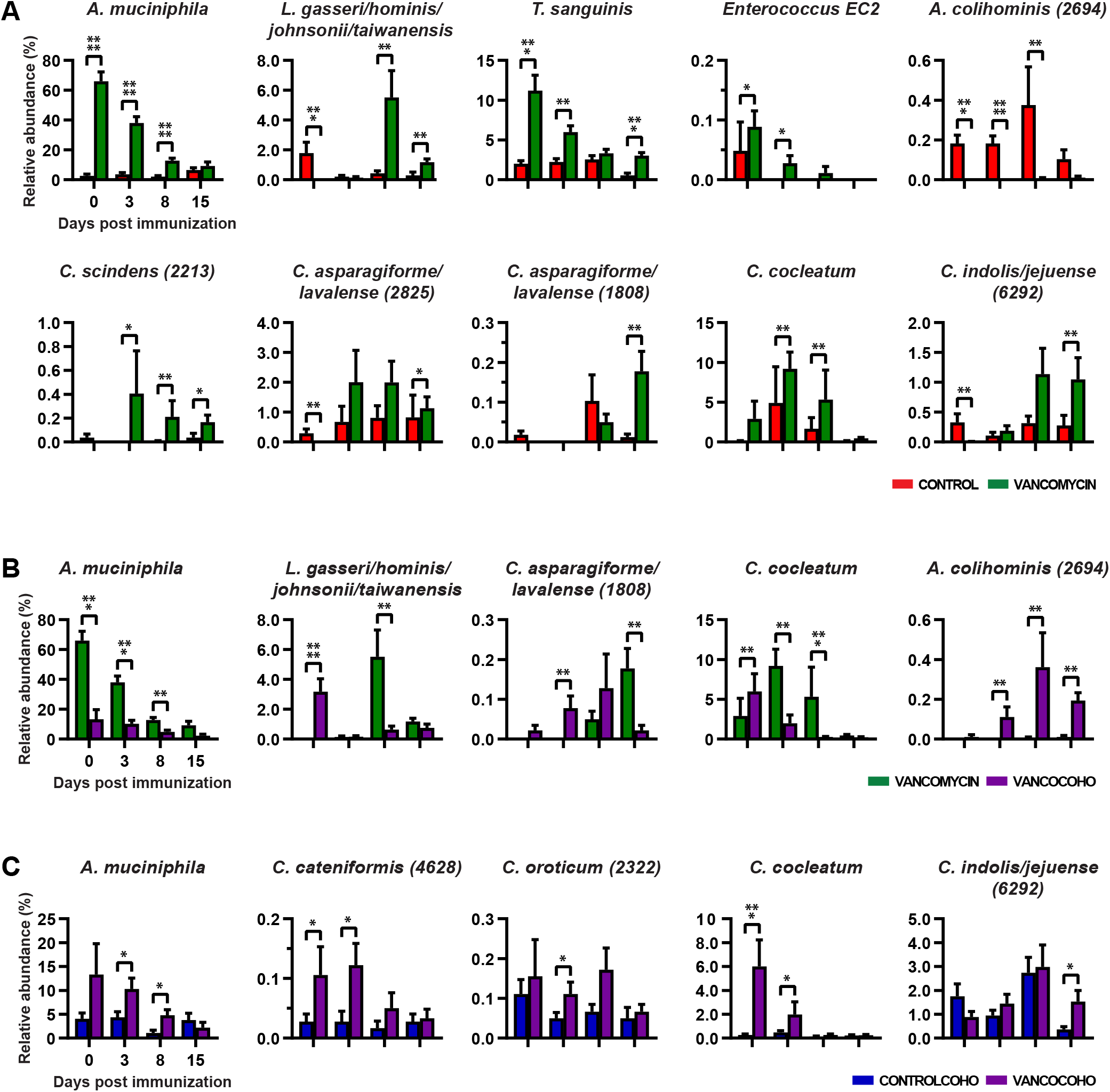
Compositional differences of selected species in untreated and vancomycin treated mice during EAE. Bar plots showing changes in the relative abundance of selected species altered at the indicated time points in: **A**. single-treatment housed untreated mice (Control) and single-treatment housed vancomycin mice (Vancomycin); **B**. single-treatment housed vancomycin mice (Vancomycin) and vancomycin treated mice cohoused with untreated mice (VancoCoho); **C**. untreated mice cohoused with vancomycin treated mice (ControlCoho) and vancomycin treated mice cohoused with untreated mice (VancoCoho). Results are presented as mean ± SEM (n = 11-12 mice/group), *p<0.05; **p<0.01; ***p<0.001; ****p<0.001. OTU = Operational taxonomic unit; Numbers in parenthesis represent OTU. EC1: *Enterococcus canintestini/canis/dispar/ durans/faecalis/faecium/hirae/ lactis/mundtii/ratti/rivorum/villorum*; EC2: *Enterococcus casseliflavus/gallinarum/saccharolyticus*.

### Correlation between bacterial abundance and EAE severity

To identify gut commensals that are associated with EAE severity, we performed Spearman correlation analysis. We identified 36 bacteria that positively correlate with EAE severity and were enriched in the control group (**Fig. 5A**). In particular, we observed that five species belonging to family *Lachnospiracea* and Clostridium cluster XIVa positively correlated with EAE severity including *C. asparagiforme /celerecrescens / lavalense / sphenoides* (OTU 911), *C. jejuense* (OTU 7388), *C. indolis* (OTU 1977), *C. aldenense/boliviensis/celerecrescens/saccharolyticum* (OTU 2224) and *C. asparagiforme/boliviensis/celerecrescens/lavalense/saccharolyticum* (OTU 121) (FDR<0.05; **Fig. 5A, 5B**). These findings are consistent with a prior study reporting a positive correlation between *Clostridium* belonging to family *Lachnospiracea* and EAE severity[80]. Another study reports a positive correlation between several OTUs belonging to Clostridium XIVa and EAE cumulative disease score[87]. We observed that genera *Roseburia* and *Anaeroplasma* positively correlated with EAE severity (**Fig. 5A-B**). These findings are consistent with prior studies reporting that *Roseburia, Ruminococcus* and *Anaeroplasma* positively correlate with EAE severity[77, 80]. We observed that the genus *Lactobacillus* correlates positively with EAE severity at 0 dpi and negatively with EAE severity at 15 dpi (**Fig. 5A-B**). These findings are consistent with prior studies reporting that some *Lactobacillus* species/strains ameliorate EAE while others exacerbate EAE[79–81, 88]. The majority of bacteria found to have a positive correlation with EAE severity have not been studied in EAE yet including *Dorea formicigenerans, Anaerosporobacter, Robinsoniella, Flavonifractor, Papillibacter, Sporobacter* and *Anaerotruncus colihominis* (**Fig. 5A**). We identified 13 bacteria that negatively correlated with EAE severity and were enriched in vancomycin treated mice (**Fig. 5A**). We found that *A. muciniphila* negatively correlate with EAE severity (**Fig. 5A-B**). Consistent with these findings, we previously reported that mice colonized with MS-derived or commercially obtained *A. muciniphila* are protected from EAE[40, 47]. Three species belonging to Clostridium sensu stricto: *C. chauvoei, C. celatum* and *C. chauvoei/quinii* (OTU 8778) negatively correlated with EAE severity (**Fig. 5A**). These findings are consistent with prior studies reporting a negative correlation between OTUs belonging to Clostridium sensu stricto and EAE cumulative disease score[87]. Several genera that negatively correlated with EAE severity have not yet been investigated in the EAE mice model including *Turicibacter sanguinis, Clostridium cocleatum, Olsenella profusa* and *Enterococcus* (**Fig. 5A**).

**Figure 5.**
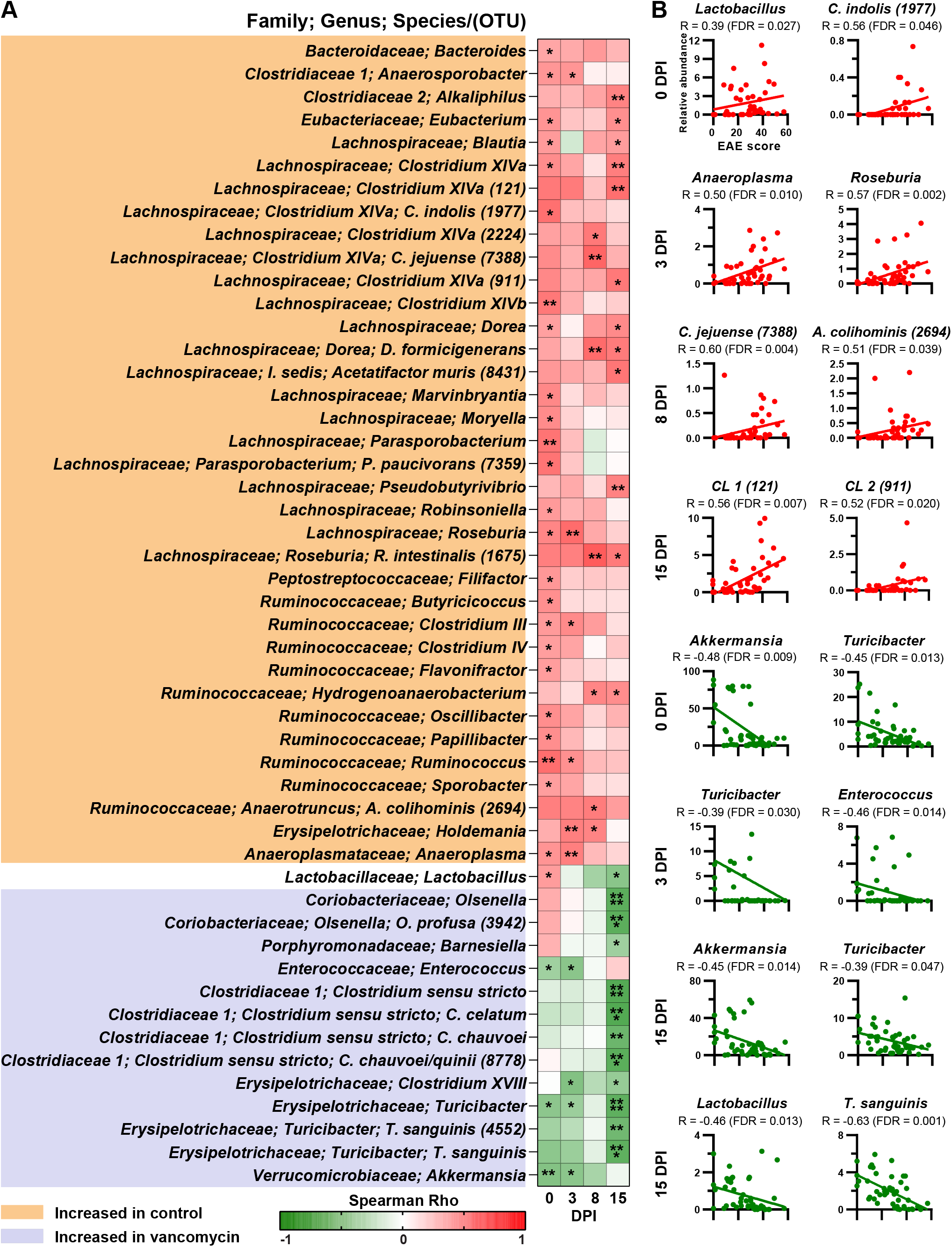
Microbiota associated with EAE severity. Spearman’s correlation between relative abundance of indicated taxa and cumulative EAE clinical score. **A**. Correlation matrix for all taxa at the lowest classifiable levels showing significant correlations at the indicated time points (FDR<0.05). *p<0.05; **p<0.01; ***p<0.001; ****p<0.001. **B**. Scatter plots of selected taxa at the lowest classifiable levels showing significant correlations at the indicated time points (FDR<0.05). OTU = operational taxonomic unit; R= correlation coefficient; Numbers in parenthesis represent OTU. CL1: *Clostridium asparagiforme/boliviensis/celerecrescens/lavalense/ saccharolyticum*. CL2: *Clostridium asparagiforme/celerecrescens/lavalense/sphenoides*.

### *A. colihominis* ameliorates EAE and induces RORγt^+^ regulatory T cells in the mesenteric lymph nodes

To this date, no human gut derived bacteria has been shown to exacerbate EAE. Given prior report that *A. colihominis* is increased in the gut of MS patients and given that it is depleted in mice protected from EAE, we hypothesized that *A. colihominis* might be an important driver of the pro-inflammatory response in EAE/MS. However, *A. colihominis* is a butyrate producing bacteria[89, 90] which could potentially play a beneficial role given prior reports that butyrate ameliorates EAE via induction of Tregs[85, 86]. Thus, whether *Anaerotruncus* contributes to disease or could protect against CNS autoimmunity needed to be tested experimentally. To investigate this, B6 mice were fed *A. colihominis, Enterococcus faecalis*, a commensal gut microbe not altered in EAE mice, or phosphate-buffered saline (PBS) 3 days per week for the entire duration of the experiment to maintain steady level of the bacteria (**Fig. 6A**). To confirm that mice were successfully colonized with *A. colihominis* and *E. faecalis*, feces were collected from mice 3 weeks post initiation of gavage, fecal DNA extracted and bacterial abundance was assessed via quantitative PCR (**Fig. 6B**). Three weeks post initiation of gavage, EAE was induced by immunizing mice with MOG. We found that mice fed *A. colihominis* had less severe disease than mice fed *E. faecalis* or PBS (**Fig. 6C**). Hence, *A. colihominis* ameliorates EAE whereas mice fed *E. faecalis* developed slightly more severe disease than PBS treated mice. These findings support a beneficial role for *A. colihominis* in EAE/MS.

**Figure 6.**
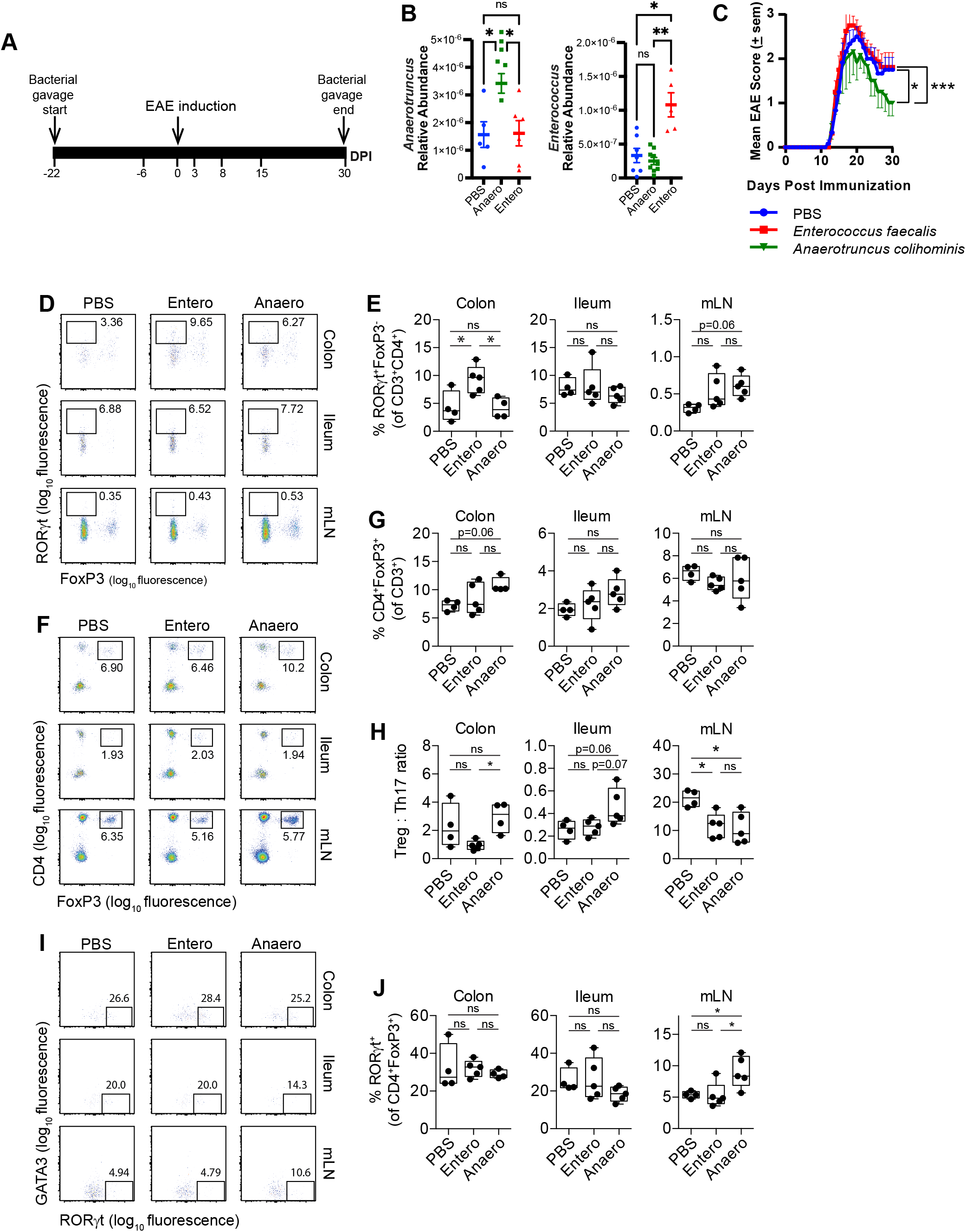
Effect of *Anaerotruncus colihominis* and *Enterococcus faecalis* on EAE development and intestinal T cell populations. **A**. Schematic design. Mice were orally gavaged daily with PBS, *Anaerotruncus colihominis* or *Enterococcus faecalis* for the entire duration of the experiment. Mice were immunized with MOG for EAE induction. **B**. qPCR quantification of the relative abundance of *Anaerotruncus colihominis* (*Anaerotruncus*) and *Enterococcus faecalis* (*Enterococcus*) by measurement of 16S rRNA gene, referenced to universal bacterial 16S rRNA gene in conventionally raised B6 mice orally gavaged with *Anaerotruncus colihominis* (Anaero), *Enterococcus faecalis* (Entero) or vehicle (PBS). Results are presented as mean ± SEM (n = 5-8). Each point represents data from one mouse. Statistical comparisons are made using one-way ANOVA and Tukey’s post hoc test. *p<0.05, **p<0.01, ns - not significant (p>0.05). **C**. Mean EAE clinical scores overtime. Results are presented as Mean ± SEM (n= 8 mice/group); the Friedman test based on scores from 15 DPI until the end of the experiment and Dunn’s multiple comparison test. *p<0.05, ***p<0.001. **(D and E)** The proportion of Th17 cells (RORγt^+^ of CD3^+^CD4^+^CD8^-^FoxP3^-^ cells) in colon and ileum lamina propria and mesenteric lymph nodes (mLN) of naïve conventionally raised mice orally gavaged with Anaero, Entero or PBS. **(F and G)** The proportion of regulatory T cells (CD4^+^FoxP3^+^ of CD3^+^ cells) in colon and ileum lamina propria and mLN of naïve conventionally raised mice orally gavaged with Anaero, Entero or PBS. **H**. The ratio of regulatory and Th17 cells in colon and ileum lamina propria and mLN of naïve conventionally raised mice orally gavaged with Anaero, Entero or PBS. **(I and J)** The proportion of RORγt^+^ regulatory T cells (RORγt^+^ of CD4^+^FoxP3^+^ cells) in colon and ileum lamina propria and mLN of naïve conventionally raised mice orally gavaged with Anaero, Entero or PBS. Boxplots show the median and interquartile range (IQR) with error bars showing the range. Each point shows data from one mouse. Statistical comparisons are made using ANOVA with Tukey correction. *p<0.05, ns – not significant (p>0.05).

To examine the effect of *A. colihominis* and *E. faecalis* on gut and peripheral immunity, naïve B6 mice were fed *A. colihominis, E. faecalis* or PBS three times per week for 3 weeks. Mice were then sacrificed and lamina propria, mesenteric lymph nodes and spleen were collected for immune profiling. We observed increased frequency of Th17 cells (CD3^+^CD4^+^RORγt^+^) in the colon of mice fed *E. faecalis* compared to mice who received PBS or *A. colihominis* (**Fig. S5A, 6D-E**). No difference in the frequency of Th17 cells (CD3^+^CD4^+^RORγt^+^) was seen in the ileum or mLN of these mice (**Fig. 6D-E**). We also observed no difference in level of IL17A, IFNγ or IL10 in the gut or spleen of these mice (**Fig. S5B-D**). GMCSF was increased in the colon of *A. colihominis* fed mice compare to *E. faecalis* or PBS treated mice but no difference in GMCSF level was observed in the ileum, mLN or spleen of these mice (**Fig. S5E**). These findings suggest that *E. faecalis* may exacerbate EAE via induction of Th17 cells in the colon. Consistent with this notion, one study reported that blocking encephalitogenic Th17 cell entry into the colon ameliorates EAE[91]. Furthermore, another study found a positive correlation between frequency of intestinal Th17 cells and disease activity in MS[57].

Next, we looked at the frequency of T regulatory cells (CD4^+^Foxp3^+^) in the gut of these mice and found no difference in the frequency of these cells in the ileum and mLN (**Fig. 6F-G**). There was a trend toward increase frequency of T regulatory cells (CD4^+^Foxp3^+^) in the colon of mice fed *A. colihominis* compared to mice fed *E. faecalis* or PBS (**Fig. 6F-G**). We observed that the Treg:Th17 ratio was trending up in the colon and ileum of B6 mice fed *A. colihominis* compare to mice fed *E. faecalis* or PBS (**Fig. 6H**). We found increased Treg:Th17 ratio in the mLN of PBS fed mice compare to mice fed *E. faecalis* and *A. colihominis* (**Fig. 6H**). A study reported that peripheral RORγt^+^ regulatory T cells suppress myelin-specific Th17 cell-mediated CNS auto-inflammation in a passive EAE model[92]. Hence, we investigated the effect of *A. colihominis* and *E. faecalis* on the frequency of peripheral RORγt^+^ regulatory T cells. We found no difference in the frequency of peripheral RORγt^+^ regulatory T cells in the spleen of these mice (**Fig. S5F**). Studies have demonstrated that intestinal RORγt^+^ Treg cells are microbiota dependent, enriched in the gut and have a strongly suppressive capacity during intestinal inflammation[93, 94]. Hence, we next examined the frequency of intestinal RORγt^+^ Treg in these mice. The frequency of RORγt^+^ Treg was comparable in all 3 groups in the colon and ileum (**Fig. 6I-J**). We observed increased frequency of RORγt^+^ regulatory T cells in the mLN of mice fed *A. colihominis* compared to mice fed *E. faecalis* or PBS (**Fig. 6I-J**). These findings are consistent with other studies reporting that a selected mixture of clostridia strains from the human microbiota including *A. colihominis* induces T regulatory cells in mouse colonic lamina propria[32, 82]. Taken together, our findings suggest that elevated *A. colihominis* in MS patients may be a host triggered response to suppress neuroinflammation. Dendritic cells from the mesenteric lymph node (mLN) are known to play a crucial role in Treg induction[95–97]. Hence, we next examined the frequency of tolerogenic dendritic cells in the mLN of these mice. We found no difference in the frequency of tolerogenic dendritic cells (CD11c^+^CD11b^+^CD103^+^) in the mLN of these mice (**Fig. S6A-C**). Hence, *A. colihominis* does not induce tolerogenic dendritic cells. Furthermore, the frequency of activated dendritic cells (CD11c^+^CD11b^+^CD80^+^) was comparable in all 3 groups in the mLN and spleen (**Fig. S6D-E**).

## Discussion

Several studies have identified alterations in the gut microbiome of MS patients and it is now accepted that the microbiome plays an important role in the pathophysiology of MS. Nevertheless, given that the mechanisms by which the microbiome affects MS have not been well defined and many confounding factors exist, findings from human microbiome studies must be validated in mice. Perturbations of the gut microbiota composition of mice have been shown to influence EAE development. Yet, few studies have characterized the microbiome during EAE and link changes in the EAE microbiome to disease severity. Hence, here we investigated changes in the gut microbiota of untreated and vancomycin treated EAE mice at multiple time points before and after EAE induction to identify gut commensals with neuro-immunomodulatory potential.

We found that vancomycin treatment ameliorates EAE and this protective effect is mediated via the microbiota. We observed an enrichment of Tregs inducing bacterial species in the gut of vancomycin treated mice including members of Clostridium cluster XIVa. We next investigated the microbiome of untreated and vancomycin treated mice to identify taxa that regulate neuroinflammation. We observed a negative correlation between *A. muciniphila* and EAE severity. Interestingly, we and others have reported increased *A. muciniphila* in MS patient compared to healthy control (HC)[11, 14, 15, 40]. In a recent study, we observed that *Akkermansia* negatively correlated with disability and T2 lesion volume and positively correlated with brain volume. These findings are consistent with our results showing that MS-derived *Akkermansia* ameliorates EAE and this protective effect is associated with decreased in RORγt^+^ and IL-17^+^ producing γδ T cells[40]. Consistent with our findings in the B6 model, another study found that mice with higher levels of *Akkermansia* had less progression in the non-obese diabetic (NOD) progressive EAE model[98]. Taken together, our findings suggest that increased Akkermansia could be a beneficial compensatory microbiome response in EAE and MS.

We also found a negative correlation between level of *Lactobacillus* and EAE severity at 15 DPI. These findings are consistent with prior studies reporting that daily administration of a mixture of *Lactobacillus* strains in EAE was effective both at preventing disease development and reversing established disease, an outcome that was IL-10 dependent and correlated with Tregs induction in mesenteric lymph nodes and the CNS[79]. Similarly, treatment with a mixture of *Lactobacillus plantarum* of human origin and *Bifidobacterium animalis* attenuated EAE clinically and induced Tregs in lymph nodes and spleen[99]. Oral administration of human derived *L. reuteri* after immunization ameliorated EAE with decreased in Th1/Th17 subsets and related cytokines[80]. In humans, one study report decreased *Lactobacillus* in MS patients[12]. We have previously reported that administering a probiotic mixture containing four *Lactobacilli* species to MS patients twice daily for 2 months leads to reduce pro-inflammatory markers in peripheral monocytes[58]. Consistent with our findings, two small double-blinded randomized controlled trials in MS patients receiving a mix of *Lactobacillus* and *Bifidobacterium* daily for 12 weeks showed significant improvements in disability score, depression, anxiety, and inflammatory markers including reduced IL-8 and TNF-alpha expression in peripheral blood mononuclear cells (PBMCs)[59, 100]. Hence these results suggest that decreased *Lactobacillus* levels in the gut of EAE mice and MS patients could promote CNS inflammation.

We observed a negative correlation between EAE severity and abundance of *Turicibacter sanguinis, Olsenella* and 3 species belonging to Clostridium sensu stricto: *C. chauvoei/quinii* (OTU 8778), *C. chauvoei* and *C. celatum*. Interestingly, we have previously reported elevated *T. sanguinis* and *Olsenella* in MS[40]. Another study reports increased Clostridium sensu stricto in MS patients but they did not identify these taxa at the species level[101]. Little is known about the functions of these bacteria in neurologic diseases. Hence, more studies are needed to investigate the role of these bacteria in EAE/MS. Several other genera that positively correlated with EAE severity have neither been implicated in MS nor have they been investigated in EAE mice including *Flavonifractor, Anaerosporobacter* and *Papillibacter*.

We found a positive correlation between EAE severity and 6 taxa belonging to Clostridium cluster XIVa including *C. indolis* (OTU 1977) and *C. jejuense* (OTU 7388). Interestingly, we have reported elevated *C. scindens* in MS patients [40]. Another study reported decreased species belonging to Clostridium cluster XIVa in MS patients but they did not identify those bacteria at the species level[41]. Several members of Clostridium cluster XIVa are butyrate producers and butyrate has been shown to induce T regulatory cells[32, 82, 83, 102]. Furthermore, several studies have reported decreased butyrate producers in MS patients[11, 41, 43, 45]. Hence, it is conceivable that loss of butyrate producers is implicated in the pathophysiology of MS and more studies are needed to identify species belonging to Clostridium cluster XIVa that play a role in MS pathophysiology.

One of the most striking differences between the microbiota of vancomycin treated mice and untreated EAE mice was depletion of *Anaerotruncus colihominis* (OTU 2694) in the single-treatment housed vancomycin group. We found a positive correlation between *A. colihominis* and EAE severity. Interestingly, we and others have previously reported increased abundance of *A. colihominis* in MS patients[19, 40]. To this date, no human derived bacteria has been shown to exacerbate EAE. Given that *A. colihominis* is depleted from the gut of mice protected from EAE, positively associated with EAE severity and is increased in MS patients, we hypothesized that it would exacerbate EAE. However, *A. colihominis* is a butyrate producer[90] and butyrate has been shown to ameliorate EAE via induction of T regs[85, 86]. Thus, whether *Anaerotruncus* contributes to disease or could protect against CNS autoimmunity needed to be tested experimentally. We found that *A. colihominis* dampened EAE severity and this was associated with increased frequency of RORγt^+^ T regulatory cells in the mesenteric lymph nodes. RORγt^+^ Tregs, secrete IL-10, a cytokine critical for immunoregulation in EAE, suggesting a possible mechanism by which these cells attenuate disease[103–105]. Taken together, our findings suggest that similar to *Akkermansia muciniphila*, increased *Anaerotruncus colihominis* in MS patients represents a protective mechanism associated with recovery from the disease and is not part of the pathogenic mechanism that induces or maintains the disease. Studies have demonstrated that intestinal RORγt^+^ Treg cells are microbiota dependent, enriched in the gut and have a strongly suppressive capacity during intestinal inflammation[93, 94]. Hence, these findings suggest that suppressing intestinal inflammation may result in decreased CNS inflammation. To this date, intestinal RORγt^+^ Treg cells have not been implicated in CNS autoimmune diseases hence, future studies should investigate the role of these Tregs in EAE mice and MS.

## Conclusions

In summary, we identified 50 gut commensals that correlate with EAE severity. Interestingly, most of the bacteria that showed a correlation with EAE severity exist in the human gut. As stated above, some of these bacteria have been implicated in MS. We also identified several bacteria that correlate with EAE severity that have not yet been implicated in MS. Some of our correlation findings have already been validated in the EAE mice model. In addition, we experimentally validated our finding of a correlation between *A. colihominis* and EAE severity. This approach can serve as a framework to test additional candidates that we identified in our study. Furthermore, future studies could perform shotgun metagenomics and metabolomics to identify species/strains specific factors in the bacteria we have identified as well as metabolites that modulate neuroinflammation in the context of EAE/MS. The findings from these studies will ultimately lead to the identification of bacteria communities that are either pathogenic or neuroprotective in MS. This in turn will facilitate the development of probiotics and other gut bacteria derived products for the prevention and treatment of MS.

## List of Abbreviations

ALS: Amyotrophic Lateral Sclerosis
B6: C57BL/6J mice strain
BHI: Brain Heart Infusion
CFA: Complete Freund’s Adjuvant
CNS: Central Nervous System
DPI: Days Post Immunization
EAE: Experimental Autoimmune Encephalomyelitis
EDSS: Expanded Disability Status Scale
FDR: False Discovery Rate
GF: Germ Free
GMCSF: Granulocyte Macrophage Stimulating Factor
HC: Healthy Control
IFNγ: Interferon gamma
IL 17: Interleukin 17
IL 10: Interleukin 10
LEfSe: Linear Discriminant Analysis Effect Size
mLN: mesenteric Lymph Node
MOG: Myelin Oligodendrocyte Glycoprotein
MS: Multiple Sclerosis
OTU: Operational Taxonomic Unit
PBS: Phosphate Buffered Saline
PBMCs: Peripheral Blood Mononuclear Cells
RORγt: RAR-related Orphan Receptor gamma T
SFB: Segmented Filamentous Bacteria
Th: T helper cell
Treg: T regulatory cell

## Declarations

### Ethics approval and consent to participate

n/a

### Consent for publication

n/a

### Availability of data and material

The microbiota 16S rRNA sequence data will be submitted to the National Center for Biotechnology Information Short Read Archives and made available to the public once the paper is published. Further information and requests for resources and reagents should be directed to and will be fulfilled by the Lead Contact, Stephanie Tankou.

### Authors’ Contributions

S.K.T, P.B, G.J.B, E.M.S and K.I performed experiments; L.M.C, P.B, D.S.W and J.C.C analyzed the microbiome data; S.L assisted with animal experimentation; G.J.B and S.K.T analyzed the FACS data; S.K.T, P.B, G.J.B and D.S.W generated the figures; S.K.T and P.B wrote the manuscript with input from all authors. S.K.T conceived the study, designed the analysis and finalized the manuscript.

## Acknowledgements

We acknowledge the support of the Dean’s Flow Cytometry CORE at the Icahn School of Medicine at Mount Sinai.

## Figure Legends

**Figure S1.**
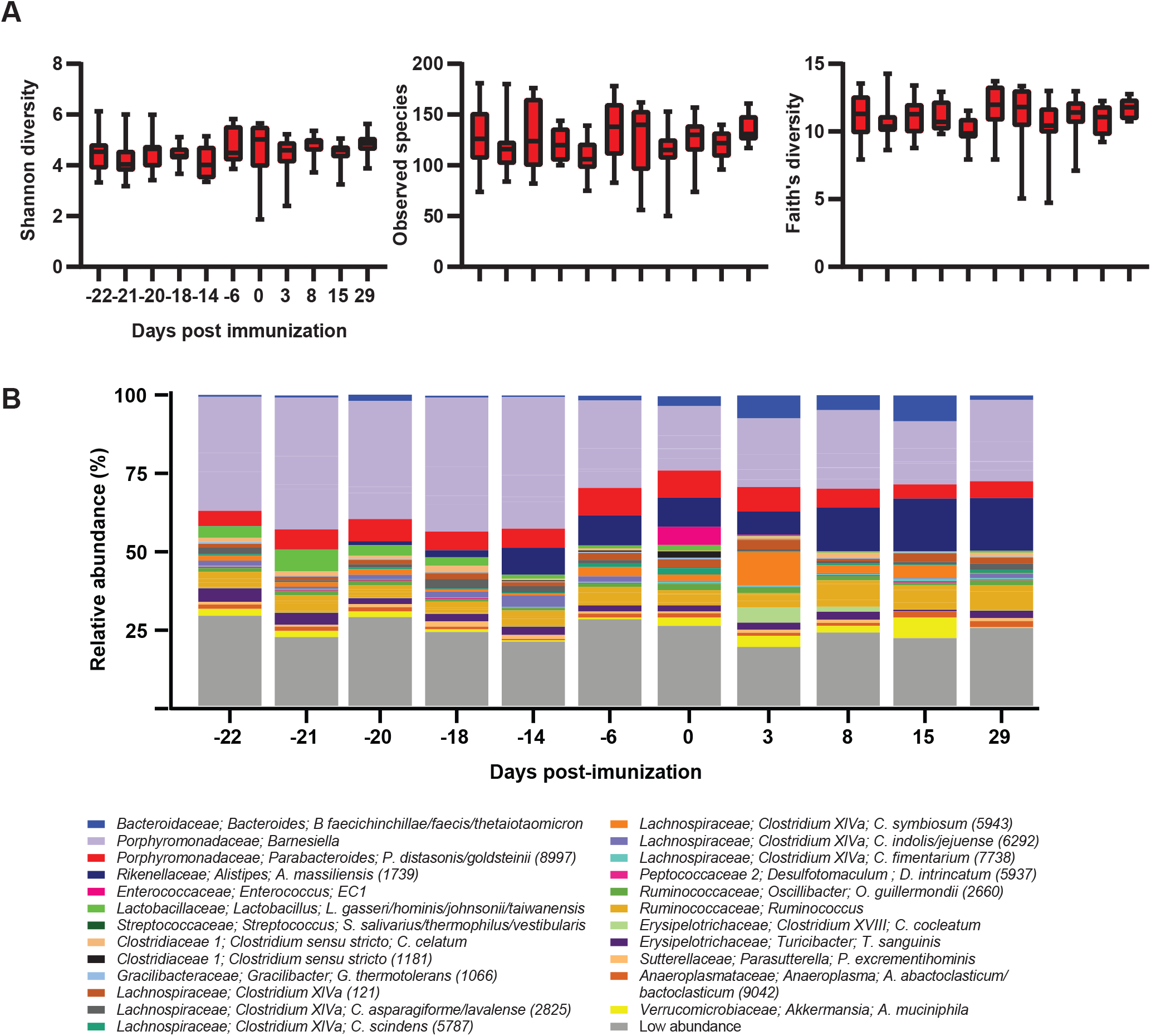
Changes in alpha diversity and microbiota composition during EAE. **A**. α-Diversity metrics for Shannon diversity, Observed species and Faith’s Diversity were calculated at an average sampling depth of 1,500 reads per sample. No significant differences were observed for any of the diversity estimators analyzed (Mixed-effect model followed by Dunnett’s test) at the indicated time points. **B**. Taxa plots showing compositional differences in fecal microbiota in mice at the indicated time points. EC1: *Enterococcus canintestini/canis/dispar/ durans/faecalis/faecium/hirae/ lactis/mundtii/ratti/rivorum/villorum*.

**Figure S2.**
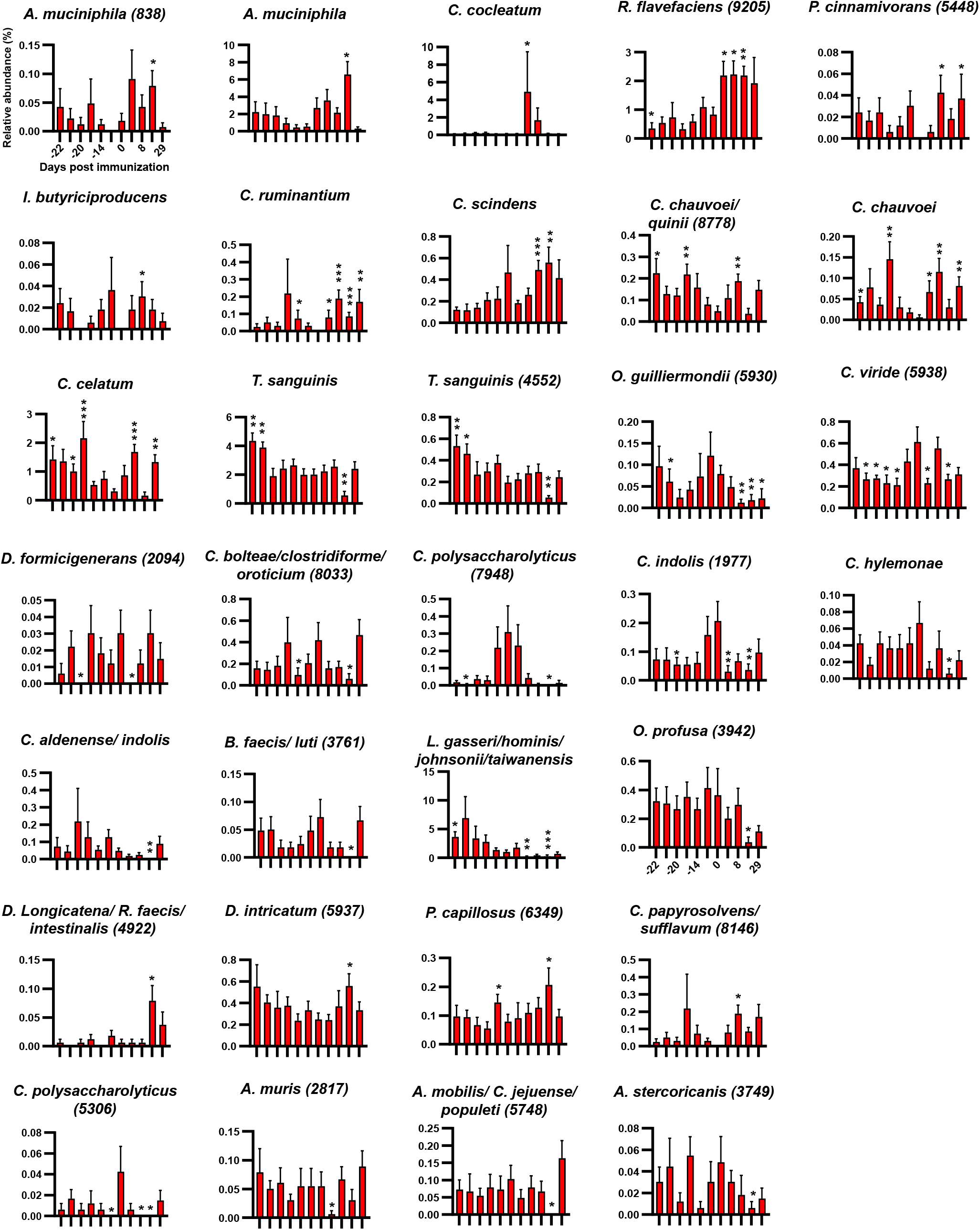
Compositional differences in the microbiota during EAE. Bar plots showing changes in the relative abundance of all significantly altered species identified by Linear Discriminant Analysis effect size. Results are presented as mean ± SEM (n = 11-12 mice). Asterisks represent significant difference at the time point indicated compare to day 0 (prior to immunization). *p<0.05; **p<0.01; ***p<0.001. Numbers in parenthesis represent operational taxonomic unit.

**Figure S3.**
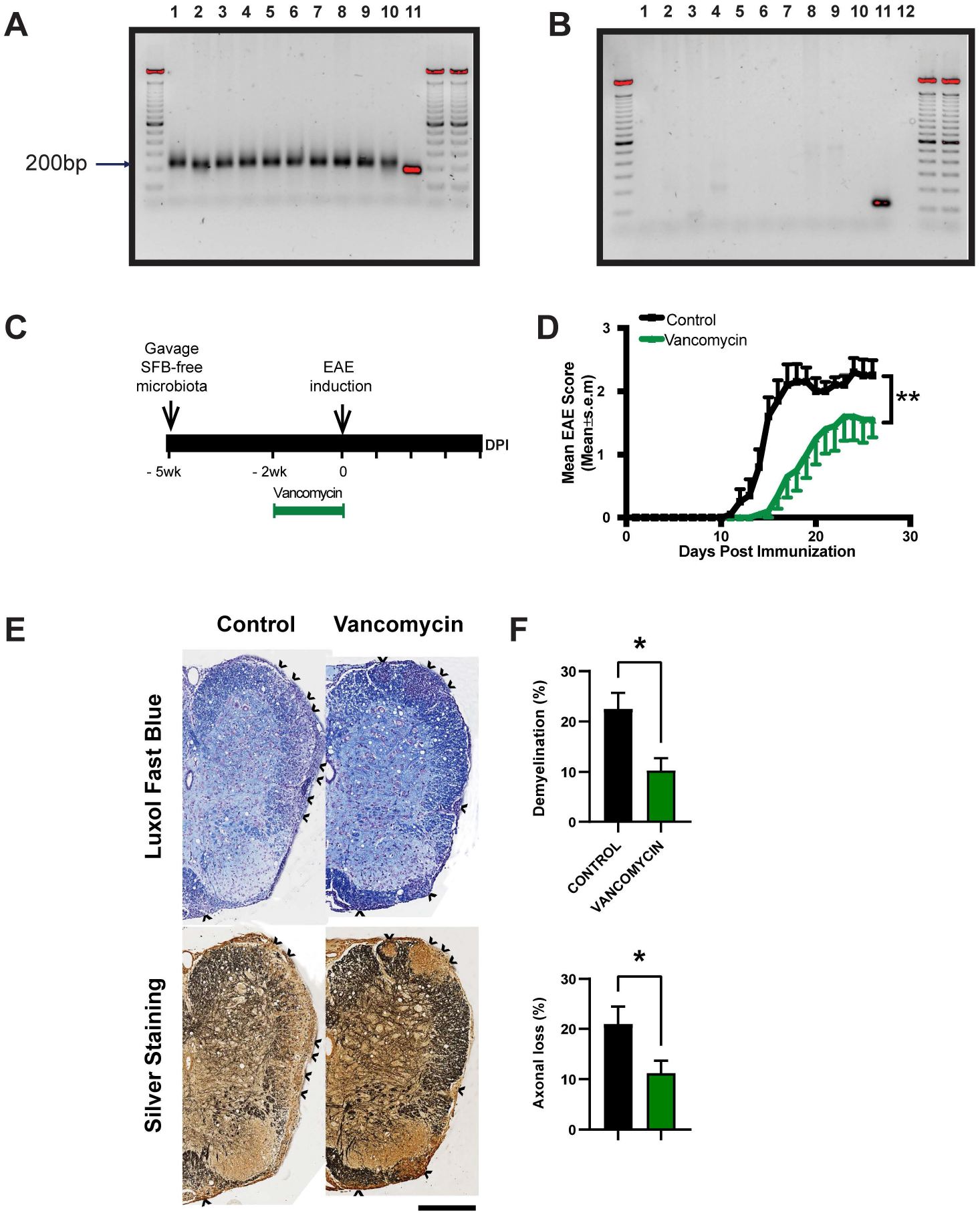
Effect of Vancomycin on EAE development in germ free mice colonized with SBF-free microbiota. DNA agarose gel showing PCR products of fecal DNA amplify with **A**. 16S rRNA gene universal primers and **B**. Segmented Filamentous Bacteria (SFB) specific primers. Lanes 1-10 represent fecal DNA from specific pathogen free B6 mice from the Jackson laboratory. Lane 11 is fecal DNA from germ-free mice monocolonized with SFB. Lane 12 of gel in panel B is empty. **C**. Experimental scheme of fecal transfer experiment in germ free mice. Germ free mice were orally gavaged with SFB-free microbiota. Three weeks post gavage, half of the colonized GF mice were on normal drinking water and the remaining half was on drinking water supplemented with vancomycin (0.5g/L) for 2 weeks. Mice were immunized with MOG for EAE induction. **D**. Mean EAE clinical scores overtime. Error bars denote Mean ± SEM (n= 10 mice/group); the Friedman test based on scores from 0 DPI until the end of the experiment and Dunn’s multiple-comparison test were performed. **p<0.01 **E**. Histopathological evaluation of demyelination with Luxol Fast Blue (LFB) and axonal loss with Bielschowsky’s silver (silver) staining. Arrows denote demyelination (LFB) and axonal loss (silver) staining of representative spinal cord sections. Scale bars, 500µm. **F**. Quantification of demyelination and axonal loss of individual mouse. Representative data of three independent experiments with n=5 mice per group are shown. Error bars denote mean ± SEM; Mann Whitney test was performed. *p<0.05.

**Figure S4.**
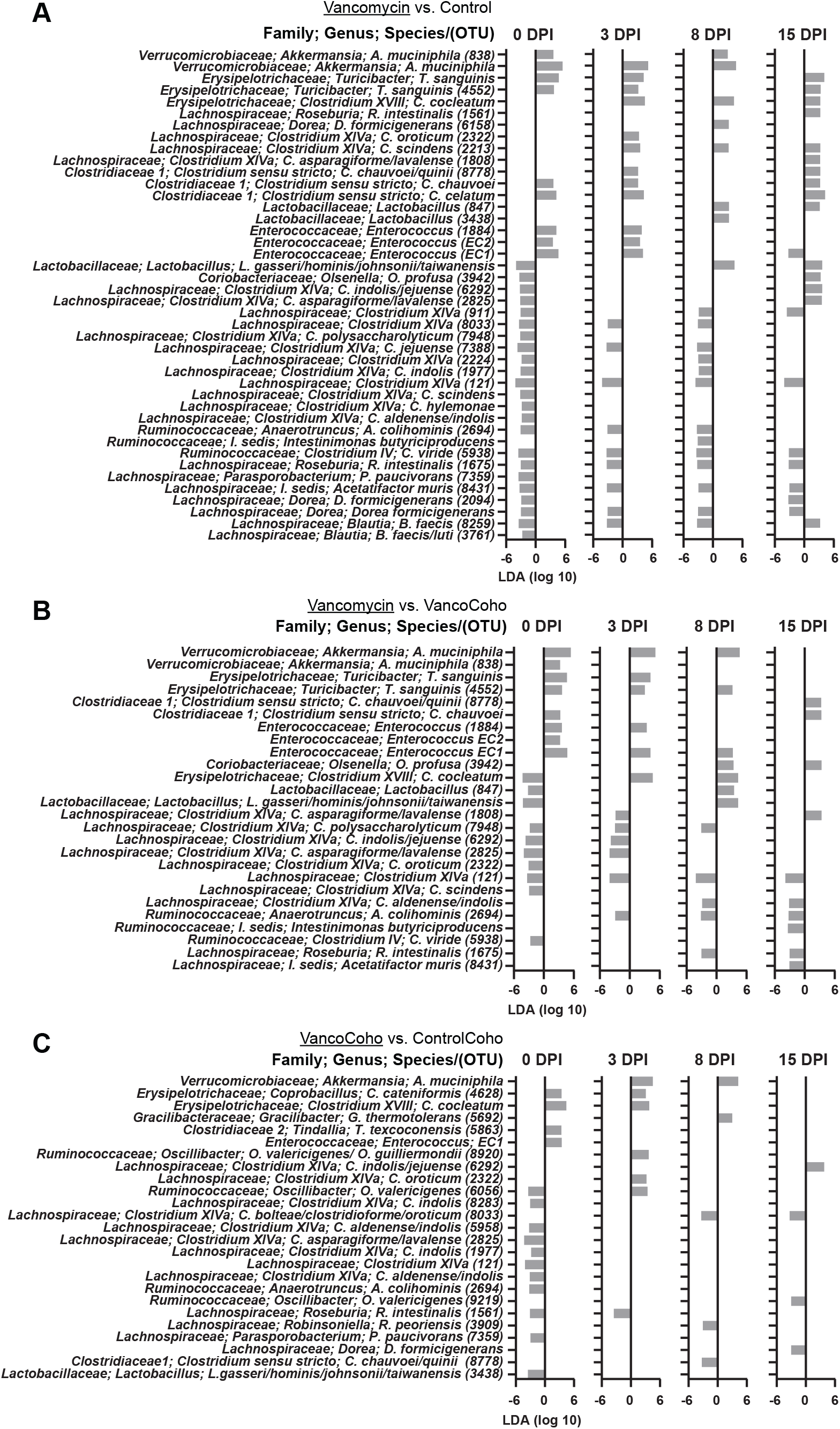
Compositional differences in the microbiota of untreated and vancomycin treated mice during EAE. Linear Discriminant Analysis (LDA) effect size of significantly altered bacteria at the species level at the indicated time points in: **A**. single-treatment housed untreated mice (Control) and single-treatment housed vancomycin mice (Vancomycin). Positive LDA effect size is increased in Vancomycin; **B**. single-treatment housed vancomycin mice (Vancomycin) and vancomycin treated mice cohoused with untreated mice (VancoCoho). Positive LDA score is increased in the underlined group; **C**. untreated mice cohoused with vancomycin treated mice (ControlCoho) and vancomycin treated mice cohoused with untreated mice (VancoCoho). Positive LDA score is increased in the underlined group. OTU = Operational taxonomic unit; Numbers in parenthesis represent OTU. EC1: *Enterococcus canintestini/canis/dispar/ durans/faecalis/faecium/hirae/ lactis/mundtii/ratti/rivorum/villorum*; EC2: *Enterococcus casseliflavus/gallinarum/saccharolyticus*.

**Figure S5.**
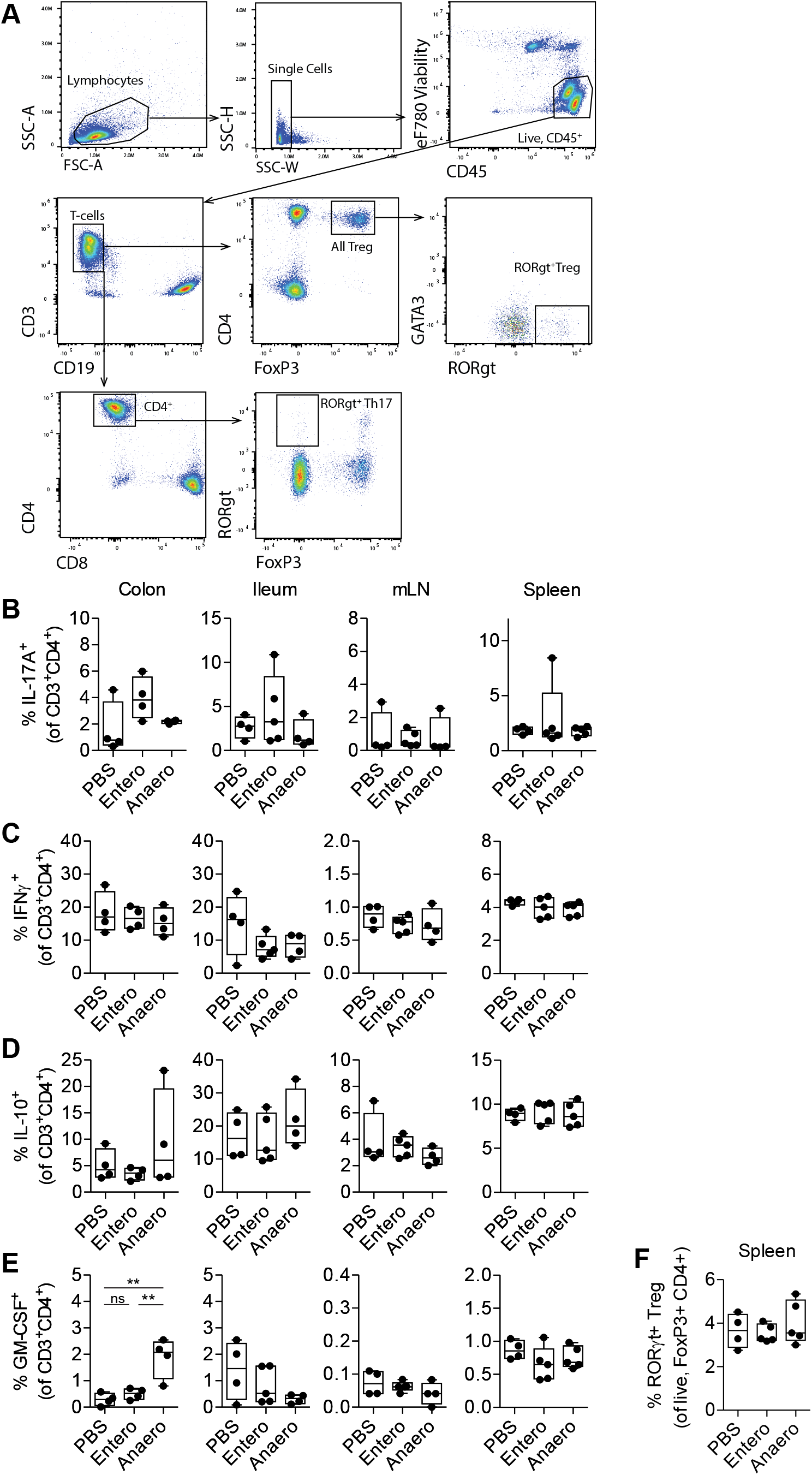
Effect of *Anaerotruncus colihominis* and *Enterococcus faecalis* on intestinal and peripheral T cell populations. **(A)** Gating strategies used to identify and quantify Th17 cells (RORγt^+^ of CD3^+^CD4^+^CD8^-^ FoxP3^-^ cells), regulatory T cells (CD4^+^FoxP3^+^ of CD3^+^ cells) and RORγt+ regulatory T cells (RORγt^+^ of CD4^+^ FoxP3^+^ cells). FSC – forward scatter, SCC – side scatter. **(B-E)** The proportion of IL-17A^+^, IFNγ^+^, IL-10^+^ and GM-CSF^+^ CD4 T cells (cytokine^+^ of CD3^+^CD4^+^ cells) in colon and ileum lamina propria, mesenteric lymph nodes (mLN) and spleen of naïve conventionally raised mice orally gavaged with *Anaerotruncus colihominis* (Anaero), *Enterococcus faecalis* (Entero) or vehicle (PBS). **F**. The proportion of RORγt^+^ regulatory T cells (RORγt^+^ of CD4^+^FoxP3^+^ cells) in the spleen of naïve conventionally raised mice orally gavaged with Anaero, Entero or PBS. Boxplots show the median and interquartile range (IQR) with error bars showing the range. Each point shows data from one mouse. Statistical comparisons are made using ANOVA with Tukey correction. **p<0.01, ns – not significant (p>0.05).

**Figure S6.**
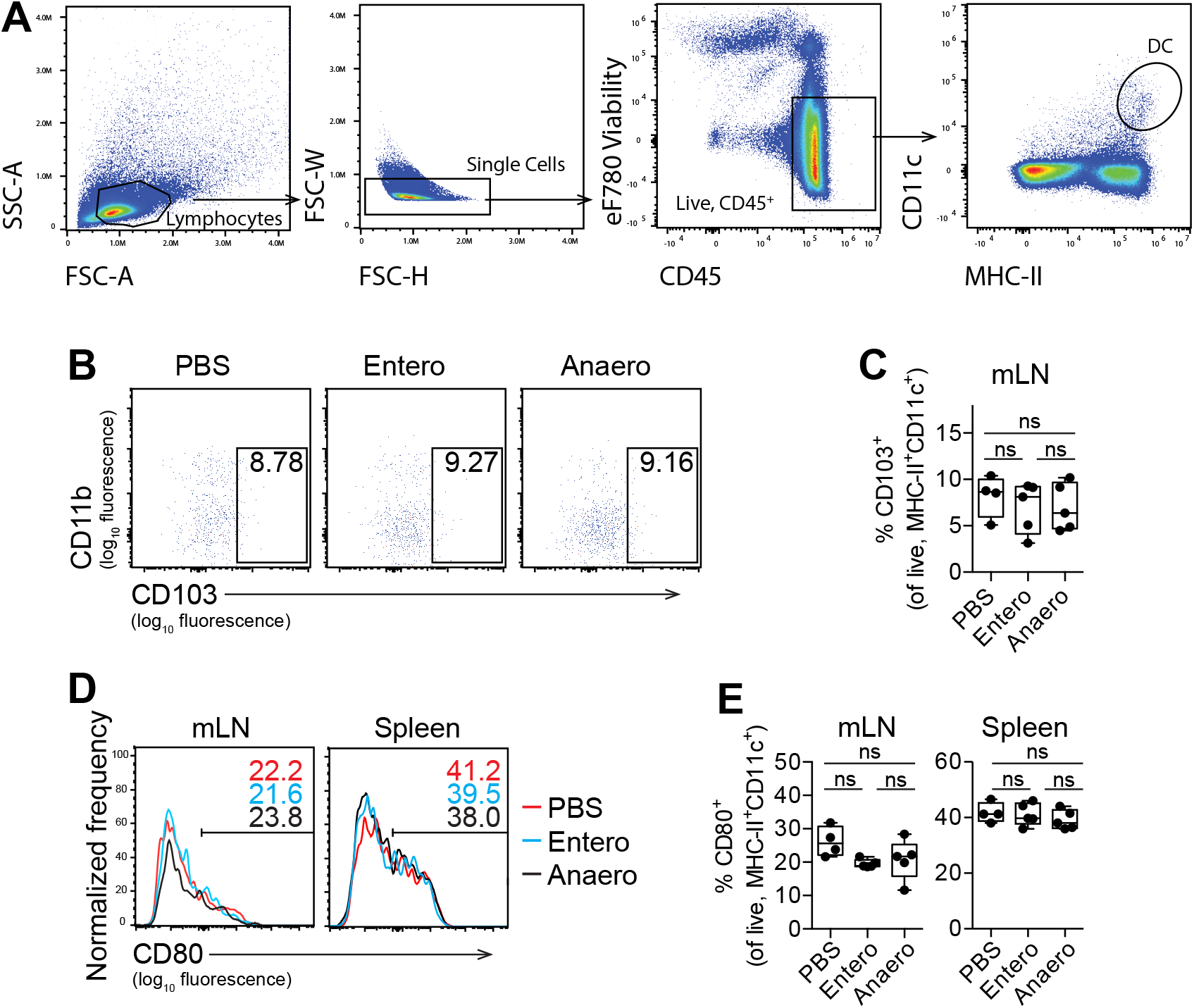
Effect of *Anaerotruncus colihominis* and *Enterococcus faecalis* on dendritic cell populations. **(A)** Gating strategies used to identify and characterize tolerogenic CD103^+^ dendritic cells (CD103^+^ of CD11c^+^MHC-II^+^ cells) and CD80^+^ dendritic cells (CD80^+^ of CD11c^+^MHC-II^+^ cells). FSC – forward scatter, SCC – side scatter. **(B and C)** The proportion of tolerogenic CD103^+^ dendritic cells (CD103^+^ of CD11c^+^MHC-II^+^ cells) in the mesenteric lymph nodes (mLN) of naïve conventionally raised mice orally gavaged with *Anaerotruncus colihominis* (Anaero), *Enterococcus faecalis* (Entero) or vehicle (PBS). **(D and E)** The proportion of CD80^+^ dendritic cells (CD80^+^ of CD11c^+^MHC-II^+^ cells) in the mLN and spleen of naïve conventionally raised mice orally gavaged with Anaero, Entero or PBS. Boxplots show the median and interquartile range (IQR) with error bars showing the range. Each point shows data from one mouse. Statistical comparisons are made using ANOVA with Tukey correction. ns – not significant (p>0.05).

## References

1. Hemmer B, Archelos JJ, Hartung H-P. New concepts in the immunopathogenesis of multiple sclerosis. Nat Rev Neurosci. 2002;3:291–301.

2. Steinman L, Martin R, Bernard C, Conlon P, Oksenberg JR. MULTIPLE SCLEROSIS: Deeper Understanding of Its Pathogenesis Reveals New Targets for Therapy*. Annu Rev Neurosci. 2002;25:491–505.

3. Prat A, Antel J. Pathogenesis of multiple sclerosis. Curr Opin Neurol. 2005;18:225–30.

4. Hafler DA, Slavik JM, Anderson DE, O’Connor KC, Jager PD, Baecher-Allan C. Multiple sclerosis. Immunol Rev. 2005;204:208–31.

5. Ristori G, Montesperelli C, Perna A, Cannoni S, Battistini L, Borsellino G, et al. Global immune disregulation in multiple sclerosis: from the adaptive response to the innate immunity. J Neuroimmunol. 2000;107:216–9.

6. Sospedra M, Martin R. IMMUNOLOGY OF MULTIPLE SCLEROSIS*. Annu Rev Immunol. 2005;23:683–747.

7. Baecher-Allan C, Kaskow BJ, Weiner HL. Multiple Sclerosis: Mechanisms and Immunotherapy. Neuron. 2018;97:742–68.

8. Frisullo G, Nociti V, Iorio R, Patanella AK, Marti A, Caggiula M, et al. IL17 and IFNγ production by peripheral blood mononuclear cells from clinically isolated syndrome to secondary progressive multiple sclerosis. Cytokine. 2008;44:22–5.

9. Olsson T, Barcellos LF, Alfredsson L. Interactions between genetic, lifestyle and environmental risk factors for multiple sclerosis. Nat Rev Neurol. 2017;13:25–36.

10. Hoogen WJ van den, Laman JD, Hart BA ‘t. Modulation of Multiple Sclerosis and Its Animal Model Experimental Autoimmune Encephalomyelitis by Food and Gut Microbiota. Front Immunol. 2017;8:1081.

11. Jangi S, Gandhi R, Cox LM, Li N, Glehn F von, Yan R, et al. Alterations of the human gut microbiome in multiple sclerosis. Nat Commun. 2016;7:12015.

12. Chen J, Chia N, Kalari KR, Yao JZ, Novotna M, Soldan MMP, et al. Multiple sclerosis patients have a distinct gut microbiota compared to healthy controls. Sci Rep-uk. 2016;6:28484.

13. Cantarel BL, Waubant E, Chehoud C, Kuczynski J, DeSantis TZ, Warrington J, et al. Gut Microbiota in Multiple Sclerosis. J Invest Med. 2015;63:729.

14. Berer K, Gerdes LA, Cekanaviciute E, Jia X, Xiao L, Xia Z, et al. Gut microbiota from multiple sclerosis patients enables spontaneous autoimmune encephalomyelitis in mice. Proc National Acad Sci. 2017;114:10719–24.

15. Cekanaviciute E, Yoo BB, Runia TF, Debelius JW, Singh S, Nelson CA, et al. Gut bacteria from multiple sclerosis patients modulate human T cells and exacerbate symptoms in mouse models. Proc National Acad Sci. 2017;114:10713–8.

16. Tremlett H, Fadrosh DW, Faruqi AA, Zhu F, Hart J, Roalstad S, et al. Gut microbiota in early pediatric multiple sclerosis: a case-control study. Eur J Neurol. 2016;23:1308–21.

17. Bhargava P, Smith MD, Mische L, Harrington EP, Fitzgerald KC, Martin KA, et al. Bile acid metabolism is altered in multiple sclerosis and supplementation ameliorates neuroinflammation. J Clin Invest. 2020. https://doi.org/10.1172/jci129401.

18. Park J, Wang Q, Wu Q, Mao-Draayer Y, Kim CH. Bidirectional regulatory potentials of short-chain fatty acids and their G-protein-coupled receptors in autoimmune neuroinflammation. Sci Rep-uk. 2019;9:8837.

19. Duscha A, Gisevius B, Hirschberg S, Yissachar N, Stangl GI, Eilers E, et al. Propionic Acid Shapes the Multiple Sclerosis Disease Course by an Immunomodulatory Mechanism. Cell. 2020. https://doi.org/10.1016/j.cell.2020.02.035.

20. Nouri M, Bredberg A, Weström B, Lavasani S. Intestinal Barrier Dysfunction Develops at the Onset of Experimental Autoimmune Encephalomyelitis, and Can Be Induced by Adoptive Transfer of Auto-Reactive T Cells. Plos One. 2014;9:e106335.

21. Mirza A, Mao-Draayer Y. The gut microbiome and microbial translocation in multiple sclerosis. Clin Immunol. 2017;183:213–24.

22. Buscarinu MC, Fornasiero A, Romano S, Ferraldeschi M, Mechelli R, Reniè R, et al. The Contribution of Gut Barrier Changes to Multiple Sclerosis Pathophysiology. Front Immunol. 2019;10:1916.

23. Wunsch M, Jabari S, Voussen B, Enders M, Srinivasan S, Cossais F, et al. The enteric nervous system is a potential autoimmune target in multiple sclerosis. Acta Neuropathol. 2017;134:281–95.

24. Belkaid Y, Hand TW. Role of the Microbiota in Immunity and Inflammation. Cell. 2014;157:121–41.

25. Geva-Zatorsky N, Sefik E, Kua L, Pasman L, Tan TG, Ortiz-Lopez A, et al. Mining the Human Gut Microbiota for Immunomodulatory Organisms. Cell. 2017;168:928–943.e11.

26. Hall JA, Bouladoux N, Sun CM, Wohlfert EA, Blank RB, Zhu Q, et al. Commensal DNA Limits Regulatory T Cell Conversion and Is a Natural Adjuvant of Intestinal Immune Responses. Immunity. 2008;29:637–49.

27. Chung H, Kasper DL. Microbiota-stimulated immune mechanisms to maintain gut homeostasis. Curr Opin Immunol. 2010;22:455–60.

28. Tanoue T, Morita S, Plichta DR, Skelly AN, Suda W, Sugiura Y, et al. A defined commensal consortium elicits CD8 T cells and anti-cancer immunity. Nature. 2019;565:600–5.

29. Ivanov II, Atarashi K, Manel N, Brodie EL, Shima T, Karaoz U, et al. Induction of Intestinal Th17 Cells by Segmented Filamentous Bacteria. Cell. 2009;139:485–98.

30. Mazmanian SK, Liu CH, Tzianabos AO, Kasper DL. An Immunomodulatory Molecule of Symbiotic Bacteria Directs Maturation of the Host Immune System. Cell. 2005;122:107–18.

31. Round JL, Lee SM, Li J, Tran G, Jabri B, Chatila TA, et al. The Toll-Like Receptor 2 Pathway Establishes Colonization by a Commensal of the Human Microbiota. Science. 2011;332:974–7.

32. Atarashi K, Tanoue T, Shima T, Imaoka A, Kuwahara T, Momose Y, et al. Induction of Colonic Regulatory T Cells by Indigenous Clostridium Species. Science. 2011;331:337–41.

33. Atarashi K, Tanoue T, Ando M, Kamada N, Nagano Y, Narushima S, et al. Th17 Cell Induction by Adhesion of Microbes to Intestinal Epithelial Cells. Cell. 2015;163:367–80.

34. Bäckhed F, Ley RE, Sonnenburg JL, Peterson DA, Gordon JI. Host-Bacterial Mutualism in the Human Intestine. Science. 2005;307:1915–20.

35. Alkanani AK, Hara N, Gottlieb PA, Ir D, Robertson CE, Wagner BD, et al. Alterations in Intestinal Microbiota Correlate With Susceptibility to Type 1 Diabetes. Diabetes. 2015;64:3510–20.

36. Kostic AD, Gevers D, Siljander H, Vatanen T, Hyötyläinen T, Hämäläinen A-M, et al. The Dynamics of the Human Infant Gut Microbiome in Development and in Progression toward Type 1 Diabetes. Cell Host Microbe. 2015;17:260–73.

37. Kostic AD, Xavier RJ, Gevers D. The Microbiome in Inflammatory Bowel Disease: Current Status and the Future Ahead. Gastroenterology. 2014;146:1489–99.

38. Scher JU, Sczesnak A, Longman RS, Segata N, Ubeda C, Bielski C, et al. Expansion of intestinal Prevotella copri correlates with enhanced susceptibility to arthritis. Elife. 2013;2:e01202.

39. Zhang X, Zhang D, Jia H, Feng Q, Wang D, Liang D, et al. The oral and gut microbiomes are perturbed in rheumatoid arthritis and partly normalized after treatment. Nat Med. 2015;21:nm.3914.

40. Cox LM, Maghzi AH, Liu S, Tankou SK, Dhang FH, Willocq V, et al. Gut Microbiome in Progressive Multiple Sclerosis. Ann Neurol. 2021;89:1195–211.

41. Miyake S, Kim S, Suda W, Oshima K, Nakamura M, Matsuoka T, et al. Dysbiosis in the Gut Microbiota of Patients with Multiple Sclerosis, with a Striking Depletion of Species Belonging to Clostridia XIVa and IV Clusters. Plos One. 2015;10:e0137429.

42. Pröbstel A-K, Baranzini SE. The Role of the Gut Microbiome in Multiple Sclerosis Risk and Progression: Towards Characterization of the “MS Microbiome.” Neurotherapeutics. 2018;15:126–34.

43. Takewaki D, Suda W, Sato W, Takayasu L, Kumar N, Kimura K, et al. Alterations of the gut ecological and functional microenvironment in different stages of multiple sclerosis. Proc National Acad Sci. 2020;117:22402–12.

44. Mirza A, Forbes JD, Zhu F, Bernstein CN, Domselaar GV, Graham M, et al. The Multiple Sclerosis Gut Microbiota: A Systematic Review. Mult Scler Relat Dis. 2019;37:101427.

45. Noto D, Miyake S. Gut dysbiosis and multiple sclerosis. Clin Immunol. 2020;:108380.

46. Volkova A, Ruggles KV. Predictive Metagenomic Analysis of Autoimmune Disease Identifies Robust Autoimmunity and Disease Specific Microbial Signatures. Front Microbiol. 2021;12:621310.

47. Liu S, Rezende RM, Moreira TG, Tankou SK, Cox LM, Wu M, et al. Oral Administration of miR-30d from Feces of MS Patients Suppresses MS-like Symptoms in Mice by Expanding Akkermansia muciniphila. Cell Host Microbe. 2019. https://doi.org/10.1016/j.chom.2019.10.008.

48. Minagar A, Alexander JS, Schwendimann RN, Kelley RE, Gonzalez-Toledo E, Jimenez JJ, et al. Combination Therapy With Interferon Beta-1a and Doxycycline in Multiple Sclerosis: An Open-Label Trial. Arch Neurol-chicago. 2008;65:199–204.

49. Mazdeh M, Mobaien AR. Efficacy of doxycycline as add-on to interferon beta-1a in treatment of multiple sclerosis. Iranian J Neurology. 2012;11:70–3.

50. Consortium T iMSMS. Household paired design reduces variance and increases power in multi-city gut microbiome study in multiple sclerosis. Mult Scler J. 2020;27:366–79.

51. Berer K, Mues M, Koutrolos M, Rasbi ZA, Boziki M, Johner C, et al. Commensal microbiota and myelin autoantigen cooperate to trigger autoimmune demyelination. Nature. 2011;479:538.

52. Lee YK, Menezes JS, Umesaki Y, Mazmanian SK. Proinflammatory T-cell responses to gut microbiota promote experimental autoimmune encephalomyelitis. Proc National Acad Sci. 2011;108 Supplement 1:4615–22.

53. Ochoa-Repáraz J, Mielcarz DW, Ditrio LE, Burroughs AR, Foureau DM, Haque-Begum S, et al. Role of Gut Commensal Microflora in the Development of Experimental Autoimmune Encephalomyelitis. J Immunol. 2009;183:6041–50.

54. Ochoa-Repáraz J, Mielcarz DW, Ditrio LE, Burroughs AR, Begum-Haque S, Dasgupta S, et al. Central Nervous System Demyelinating Disease Protection by the Human Commensal Bacteroides fragilis Depends on Polysaccharide A Expression. J Immunol. 2010;185:4101–8.

55. Shahi SK, Freedman SN, Murra AC, Zarei K, Sompallae R, Gibson-Corley KN, et al. Prevotella histicola, A Human Gut Commensal, Is as Potent as COPAXONE® in an Animal Model of Multiple Sclerosis. Front Immunol. 2019;10:462.

56. Mangalam A, Shahi SK, Luckey D, Karau M, Marietta E, Luo N, et al. Human Gut-Derived Commensal Bacteria Suppress CNS Inflammatory and Demyelinating Disease. Cell Reports. 2017;20:1269–77.

57. Cosorich I, Dalla-Costa G, Sorini C, Ferrarese R, Messina MJ, Dolpady J, et al. High frequency of intestinal TH17 cells correlates with microbiota alterations and disease activity in multiple sclerosis. Sci Adv. 2017;3:e1700492.

58. Tankou SK, Regev K, Healy BC, Tjon E, Laghi L, Cox LM, et al. A probiotic modulates the microbiome and immunity in multiple sclerosis. Ann Neurol. 2018;83:1147–61.

59. Kouchaki E, Tamtaji OR, Salami M, Bahmani F, Kakhaki RD, Akbari E, et al. Clinical and metabolic response to probiotic supplementation in patients with multiple sclerosis: A randomized, double-blind, placebo-controlled trial. Clin Nutr. 2017;36.

60. Mayo L, Trauger SA, Blain M, Nadeau M, Patel B, Alvarez JI, et al. Regulation of astrocyte activation by glycolipids drives chronic CNS inflammation. Nat Med. 2014;20:1147–56.

61. Cox LM, Sohn J, Tyrrell KL, Citron DM, Lawson PA, Patel NB, et al. Description of two novel members of the family Erysipelotrichaceae: Ileibacteriumvalens gen. nov., sp. nov. and Dubosiella newyorkensis, gen. nov., sp. nov., from the murine intestine, and emendation to the description of Faecalibacterium rodentium. Int J Syst Evol Micr. 2017;67:1247–54.

62. Caporaso JG, Lauber CL, Walters WA, Berg-Lyons D, Huntley J, Fierer N, et al. Ultra-high-throughput microbial community analysis on the Illumina HiSeq and MiSeq platforms. Isme J. 2012;6:1621.

63. Shaikh FY, White JR, Gills JJ, Hakozaki T, Richard C, Routy B, et al. A Uniform Computational Approach Improved on Existing Pipelines to Reveal Microbiome Biomarkers of Nonresponse to Immune Checkpoint InhibitorsSingle Pipeline Reanalysis. Clin Cancer Res. 2021;27:2571–83.

64. Daquigan N, Seekatz AM, Greathouse KL, Young VB, White JR. High-resolution profiling of the gut microbiome reveals the extent of Clostridium difficile burden. Npj Biofilms Microbiomes. 2017;3:35.

65. Barman M, Unold D, Shifley K, Amir E, Hung K, Bos N, et al. Enteric Salmonellosis Disrupts the Microbial Ecology of the Murine Gastrointestinal Tract▿. Infect Immun. 2008;76:907–15.

66. Grazul H, Kanda LL, Gondek D. Impact of probiotic supplements on microbiome diversity following antibiotic treatment of mice. Gut Microbes. 2016;7:101–14.

67. Ke D, Picard FJ, Martineau F, Ménard C, Roy PH, Ouellette M, et al. Development of a PCR Assay for Rapid Detection of Enterococci. J Clin Microbiol. 1999;37:3497–503.

68. Schmittgen TD, Livak KJ. Analyzing real-time PCR data by the comparative CT method. Nat Protoc. 2008;3:1101–8.

69. Britton GJ, Contijoch EJ, Mogno I, Vennaro OH, Llewellyn SR, Ng R, et al. Microbiotas from Humans with Inflammatory Bowel Disease Alter the Balance of Gut Th17 and RORγt+ Regulatory T Cells and Exacerbate Colitis in Mice. Immunity. 2019;50:212–224.e4.

70. Baker D, Lidster K, Sottomayor A, Amor S. Two Years Later: Journals Are Not Yet Enforcing the ARRIVE Guidelines on Reporting Standards for Pre-Clinical Animal Studies. Plos Biol. 2014;12:e1001756.

71. Moore SM, Khalaj AJ, Kumar S, Winchester Z, Yoon J, Yoo T, et al. Multiple functional therapeutic effects of the estrogen receptor β agonist indazole-Cl in a mouse model of multiple sclerosis. Proc National Acad Sci. 2014;111:18061–6.

72. Ponomarev ED, Veremeyko T, Barteneva N, Krichevsky AM, Weiner HL. MicroRNA-124 promotes microglia quiescence and suppresses EAE by deactivating macrophages via the C/EBP-α–PU.1 pathway. Nat Med. 2011;17:64–70.

73. Cignarella F, Cantoni C, Ghezzi L, Salter A, Dorsett Y, Chen L, et al. Intermittent Fasting Confers Protection in CNS Autoimmunity by Altering the Gut Microbiota. Cell Metab. 2018;27:1222–1235.e6.

74. Rothhammer V, Borucki DM, Tjon EC, Takenaka MC, Chao C-C, Ardura-Fabregat A, et al. Microglial control of astrocytes in response to microbial metabolites. Nature. 2018;557:724–8.

75. Muramatsu R, Kubo T, Mori M, Nakamura Y, Fujita Y, Akutsu T, et al. RGMa modulates T cell responses and is involved in autoimmune encephalomyelitis. Nat Med. 2011;17:488–94.

76. Gandy KAO, Zhang J, Nagarkatti P, Nagarkatti M. The role of gut microbiota in shaping the relapse-remitting and chronic-progressive forms of multiple sclerosis in mouse models. Sci Rep-uk. 2019;9:6923.

77. Johanson DM, Goertz JE, Marin IA, Costello J, Overall CC, Gaultier A. Experimental autoimmune encephalomyelitis is associated with changes of the microbiota composition in the gastrointestinal tract. Sci Rep-uk. 2020;10:15183.

78. Gonzalez CG, Tankou SK, Cox LM, Casavant EP, Weiner HL, Elias JE. Latent-period stool proteomic assay of multiple sclerosis model indicates protective capacity of host-expressed protease inhibitors. Sci Rep-uk. 2019;9:12460.

79. Lavasani S, Dzhambazov B, Nouri M, Fåk F, Buske S, Molin G, et al. A Novel Probiotic Mixture Exerts a Therapeutic Effect on Experimental Autoimmune Encephalomyelitis Mediated by IL-10 Producing Regulatory T Cells. Plos One. 2010;5:e9009.

80. He B, Hoang TK, Tian X, Taylor CM, Blanchard E, Luo M, et al. Lactobacillus reuteri Reduces the Severity of Experimental Autoimmune Encephalomyelitis in Mice by Modulating Gut Microbiota. Front Immunol. 2019;10:385.

81. Yamashita M, Ukibe K, Matsubara Y, Hosoya T, Sakai F, Kon S, et al. Lactobacillus helveticus SBT2171 Attenuates Experimental Autoimmune Encephalomyelitis in Mice. Front Microbiol. 2018;8:2596.

82. Atarashi K, Tanoue T, Oshima K, Suda W, Nagano Y, Nishikawa H, et al. Treg induction by a rationally selected mixture of Clostridia strains from the human microbiota. Nature. 2013;500:232.

83. Honda K, Littman DR. The microbiota in adaptive immune homeostasis and disease. Nature. 2016;535:75–84.

84. Kageyama A, Benno Y. Coprobacillus catenaformis Gen. Nov., Sp. Nov., a New Genus and Species Isolated from Human Feces. Microbiol Immunol. 2000;44:23–8.

85. Mizuno M, Noto D, Kaga N, Chiba A, Miyake S. The dual role of short fatty acid chains in the pathogenesis of autoimmune disease models. Plos One. 2017;12:e0173032.

86. Haghikia A, Jörg S, Duscha A, Berg J, Manzel A, Waschbisch A, et al. Dietary Fatty Acids Directly Impact Central Nervous System Autoimmunity via the Small Intestine. Immunity. 2015;43:817–29.

87. Montgomery TL, Künstner A, Kennedy JJ, Fang Q, Asarian L, Culp-Hill R, et al. Interactions between host genetics and gut microbiota determine susceptibility to CNS autoimmunity. Proc National Acad Sci. 2020;117:27516–27.

88. Miyauchi E, Kim S-W, Suda W, Kawasumi M, Onawa S, Taguchi-Atarashi N, et al. Gut microorganisms act together to exacerbate inflammation in spinal cords. Nature. 2020;585:102–6.

89. Lawson PA, Song Y, Liu C, Molitoris DR, Vaisanen M-L, Collins MD, et al. Anaerotruncus colihominis gen. nov., sp. nov., from human faeces. Int J Syst Evol Micr. 2004;54:413–7.

90. Narushima S, Sugiura Y, Oshima K, Atarashi K, Hattori M, Suematsu M, et al. Characterization of the 17 strains of regulatory T cell-inducing human-derived Clostridia. Gut Microbes. 2014;5:333–9.

91. Duc D, Vigne S, Bernier-Latmani J, Yersin Y, Ruiz F, GaÏa N, et al. Disrupting Myelin-Specific Th17 Cell Gut Homing Confers Protection in an Adoptive Transfer Experimental Autoimmune Encephalomyelitis. Cell Reports. 2019;29:378–390.e4.

92. Kim B-S, Lu H, Ichiyama K, Chen X, Zhang Y-B, Mistry NA, et al. Generation of RORγt+ Antigen-Specific T Regulatory 17 Cells from Foxp3+ Precursors in Autoimmunity. Cell Reports. 2017;21:195–207.

93. Sefik E, Geva-Zatorsky N, Oh S, Konnikova L, Zemmour D, McGuire AM, et al. Individual intestinal symbionts induce a distinct population of RORγ+ regulatory T cells. Science. 2015;349:993–7.

94. Yang B-H, Hagemann S, Mamareli P, Lauer U, Hoffmann U, Beckstette M, et al. Foxp3+ T cells expressing RORγt represent a stable regulatory T-cell effector lineage with enhanced suppressive capacity during intestinal inflammation. Mucosal Immunol. 2016;9:444–57.

95. Cording S, Wahl B, Kulkarni D, Chopra H, Pezoldt J, Buettner M, et al. The intestinal micro-environment imprints stromal cells to promote efficient Treg induction in gut-draining lymph nodes. Mucosal Immunol. 2014;7:359–68.

96. Coombes JL, Siddiqui KRR, Arancibia-Cárcamo CV, Hall J, Sun C-M, Belkaid Y, et al. A functionally specialized population of mucosal CD103+ DCs induces Foxp3+ regulatory T cells via a TGF-β–and retinoic acid–dependent mechanism. J Exp Medicine. 2007;204:1757–64.

97. Pezoldt J, Pasztoi M, Zou M, Wiechers C, Beckstette M, Thierry GR, et al. Neonatally imprinted stromal cell subsets induce tolerogenic dendritic cells in mesenteric lymph nodes. Nat Commun. 2018;9:3903.

98. Colpitts SL, Kasper EJ, Keever A, Liljenberg C, Kirby T, Magori K, et al. A bidirectional association between the gut microbiota and CNS disease in a biphasic murine model of multiple sclerosis. Gut Microbes. 2017;8:561–73.

99. Salehipour Z, Haghmorad D, Sankian M, Rastin M, Nosratabadi R, Dallal MMS, et al. Bifidobacterium animalis in combination with human origin of Lactobacillus plantarum ameliorate neuroinflammation in experimental model of multiple sclerosis by altering CD4+ T cell subset balance. Biomed Pharmacother. 2017;95:1535–48.

100. Tamtaji OR, Kouchaki E, Salami M, Aghadavod E, Akbari E, Tajabadi-Ebrahimi M, et al. The Effects of Probiotic Supplementation on Gene Expression Related to Inflammation, Insulin, and Lipids in Patients With Multiple Sclerosis: A Randomized, Double-Blind, Placebo-Controlled Trial. J Am Coll Nutr. 2017;:1–6.

101. Forbes JD, Chen C, Knox NC, Marrie R-A, El-Gabalawy H, Kievit T de, et al. A comparative study of the gut microbiota in immune-mediated inflammatory diseases—does a common dysbiosis exist? Microbiome. 2018;6:221.

102. Arpaia N, Campbell C, Fan X, Dikiy S, Veeken J van der, deRoos P, et al. Metabolites produced by commensal bacteria promote peripheral regulatory T-cell generation. Nature. 2013;504:451.

103. Lochner M, Peduto L, Cherrier M, Sawa S, Langa F, Varona R, et al. In vivo equilibrium of proinflammatory IL-17+ and regulatory IL-10+ Foxp3+ RORγt+ T cells. J Exp Medicine. 2008;205:1381–93.

104. Xu M, Pokrovskii M, Ding Y, Yi R, Au C, Harrison OJ, et al. c-MAF-dependent regulatory T cells mediate immunological tolerance to a gut pathobiont. Nature. 2018;554:373–7.

105. Bettelli E, Das MP, Howard ED, Weiner HL, Sobel RA, Kuchroo VK. IL-10 is critical in the regulation of autoimmune encephalomyelitis as demonstrated by studies of IL-10- and IL-4-deficient and transgenic mice. J Immunol Baltim Md 1950. 1998;161:3299–306.

